# Drivers of amphibian population dynamics and asynchrony at local and continental scales

**DOI:** 10.1101/592683

**Authors:** Hugo Cayuela, Richard A. Griffiths, Nurul Zakaria, Jan W. Arntzen, Pauline Priol, Jean-Paul Léna, Aurélien Besnard, Pierre Joly

## Abstract

1. Identifying the drivers of population fluctuations in spatially distinct populations remains a significant challenge for ecologists. Whereas regional climatic factors may generate population synchrony (i.e., Moran effect), local factors including the level of density-dependence may reduce the level of synchrony. Although divergences in the scaling of population synchrony and spatial environmental variation have been observed, the regulatory factors that underlie such mismatches are poorly understood.
2. No previous studies have investigated how density-dependent processes and population-specific responses to weather variation influence spatial synchrony at both local and continental scales. We addressed this issue in a pond-breeding amphibian, the great crested newt (*Triturus cristatus*). We used capture-recapture data collected through long-term surveys in five *T. cristatus* populations in Western Europe.
3. We found a low level of demographic synchrony at both local and continental levels. Weather has weak and spatially variable effects on survival, recruitment and population growth rate. In contrast, density-dependence was a common phenomenon (at least for population growth) in almost all populations and subpopulations.
4. Our findings support the idea that Moran’s effect is low in species where the population dynamics more closely depends on local factors (e.g. population density and habitat characteristics) than on large-scale environmental fluctuation (e.g. regional climatic variation). Those demographic feature likely have far-reaching consequences for the long-term viability of the spatially structured populations and their ability to response to large-scale climatic anomalies.

## INTRODUCTION

Understanding the mechanisms driving population dynamics is a critical challenge in ecology and conservation biology. During recent decades, researchers have paid particular attention to the mechanisms involved in the synchronization of population dynamics at different scales (Ranta et al. 1995, Bjørnstad et al. 1999, Liebhold et al. 2004, Koenig & Liebhold 2016). Three mechanisms have been widely recognized to cause population synchrony: dispersal between populations (Kendall et al. 2000, Ylikarjula et al. 2000); trophic interactions with other species that are themselves either synchronized or mobile (Ims & Andreassen 2000, Korpimäki et al. 2005); and synchronous stochastic environmental effects, called Moran’s effect (Moran 1953, Koenig 2002). While the first two factors usually drive population synchrony at relatively local spatial scales, the Moran effect is considered a major mechanism for generating population synchrony at continental level (Koenig 2002).

Moran’s (1953) concept states that synchrony in the dynamics of populations regulated by the same density-dependent processes will reflect a correlation in the environmental perturbations. Consequently, population synchrony is expected to decay with increasing distance between populations in a manner similar to that of the synchrony of a potential underlying environmental factor. However, the scaling of the population synchrony is far less than the spatial scale of variation in environmental variables (Koenig 2001, Peltonen et al. 2002, Trenham et al. 2003). For example, simulation studies have revealed that the degree of population synchrony is influenced by the form of density regulation. Specifically, population-specific density dependence and nonlinear relationships between demographic rates and density typically reduce the synchronizing effect of spatially correlated environmental noise (Ranta et al. 1995, Bjørnstad et al. 1999, Royama 2005). Secondly, theoretical models predict that spatial variation of the demographic parameters contributing to population growth rate negates the Moran effect and then reduces population synchrony (Engen & Sæther 2005, Hugueny 2006). For instance, variation in survival and recruitment caused by local environmental factors such as predation, interspecific competition and habitat quality could decrease population synchrony. Simulations have also demonstrated that the contribution of a covariate to spatial synchrony strongly depends on spatial heterogeneity in the covariate or on its effect on local dynamics (Engen & Sæther 2005, Hugueny 2006). Especially, local expression of large-scale climatic phenomena could lead to context-dependent demographic responses to weather variation (Selwood et al. 2015), which could then result in low spatial synchrony.

Several empirical studies have reported divergences in the scaling of population synchrony and spatial environmental variation (Koenig 2001, Peltonen et al. 2002, Trenham et al. 2003). However, few have empirically examined the factors that could underlie such mismatches. No previous work has investigated how population-specific density dependent processes and population-specific responses to environmental variation (e.g. weather) could affect spatial synchrony at both local and continental scales. Pond-breeding amphibians are excellent biological models to address this issue. First, population synchrony has already been highlighted in these organisms (Trenham et al. 2003) and spatially correlated environmental weather variables (i.e. temperature and precipitation) could have strong synchronizing effects. Yet, several factors with potentially desynchronizing effects also play an important role in amphibian demography. Variation in pond characteristics may dramatically affect demographic rates (Unglaub et al. 2015, 2018). Furthermore, although amphibians may be highly sensitive to climate, their demographic responses to weather variation seems highly context-dependent (Cayuela et al. 2016a, Muths et al. 2017). Moreover, density-dependence has been often reported as a key driver of amphibian population dynamics (Altwegg & Reyer 2003, Harper & Semlitsch 2007).

In this study, we examined population synchrony, demographic responses to weather and density dependence in the great crested newt (*Triturus cristatus*). We used capture-recapture data collected through long-term surveys in five *T. cristatus* populations in Western Europe over a 22-year period (1995-2016). Two of these five populations (i.e. POP1 and POP5) were spatially structured (i.e. composed of subpopulations occupying different ponds or group of ponds). This nested design allowed us to examine demographic processes at both local (i.e. within population) and continental (between population, over western Europe) levels. We predicted a relatively low level of spatial synchrony at both local and continental levels due to the characteristics of amphibian demography stated above. Second, we examined how local demographic responses to climate and density-dependence may trigger population asynchrony. In particular, we investigated how weather fluctuation and population density may affect survival, recruitment and population growth rate in the five populations and in the four subpopulations of POP1 and POP5. According to the conclusion of previous studies (Cayuela et al. 2016a, Muths et al. 2017), we expected heterogeneous demographic responses (survival, recruitment, and population growth) to temperature and rainfall within and between populations. In addition, we predicted negative density-dependence to be a common phenomenon. At the continental level, we assumed that such density-dependent processes would have a desynchronizing effect since populations have different histories, are located in different geographical areas and are not connected by dispersing individuals. At the local level, we also predicted a desynchronizing effect of density as dispersal rates are low between subpopulations (see Material and method section) and because ponds may differ in terms of local characteristics.

## MATERIAL AND METHODS

### Studied populations, capture-recapture data and weather variables

The study was conducted on five populations in France and the United Kingdom using the capture– recapture method. Two populations (POP1 and POP2) are located in southeastern England and are 2.5 km apart and separated from each other by dispersal barriers, such as roads and unsuitable habitat (Zakaria 2018). Another population (POP3) is located in western France. The last two populations (POP4 and POP5) are located in southeastern France. The population POP2, POP3 and POP4 occupied a single site for reproduction. POP1 and POP5 are spatially structured populations, both composed of four distinct subpopulations. In POP1, the four subpopulations occupy four distinct ponds, separated from each other by distances ranging from 200 to 800 m. According to Griffiths et al. (2010), there is a low level of adult movement among subpopulations with dispersal mainly occurring during the subadult phase. In POP5, the four subpopulations occupy four distinct pond groups. Each group was composed of three very close (15-30 m) ponds between which annual dispersal rates are high (<0.20; Cayuela et al. 2018a). The pond groups were separated from each other’s by a distance ranging from 60 to 430 m; the number of breeding dispersal events among ponds was low (i.e. 12 out of 2282 individuals captured during the period 1996-2015), resulting in mean dispersal rate < 0.01. These 12 individuals were discarded from our analyses.

Newts were surveyed over periods ranging from 8 to 20 years between 1995 and 2016. The newts were captured using bottle traps (POP1 and POP2, see Griffiths et al. 2010), funnel traps (POP3), dipnets (POP4, see Cayuela et al. 2017) and seine net (POP5, see Cayuela et al. 2017). The number of capture sessions performed each year varied from 1 (POP5) to 33 (POP1) during the newt activity period in ponds. Note that in capture-recapture analyses, we merged the observations of intra-annual sessions, considering one single session per year. Newts were individually identified using pit-tags in POP3 and POP5, and by photographs of belly pattern markings in POP1, POP2 and POP4. Providing the image quality is high and the identification is done by trained personnel, belly patterns are a highly reliable method for identifying individual newts (Zakaria, 2018). We recorded for each individual its maturity status and gender. Gender was assessed on the basis of the presence of a swollen cloaca and a large crest on the back in males (Hedlund 1990). As the number of juveniles was low in our datasets (juveniles only occasionally found in the ponds), the following analyses were restricted to adults. Detailed information about the capture–recapture surveys (samples size, the number of captured newts, etc.) are provided in Appendix 1.

In Western Europe, both precipitation and temperatures exhibit high synchrony declining with distance; the spatial autocorrelation remains statistically significant over a very large distance (up to 2500 km; Koenig 2002). We thus assumed that these two meteorological parameters could act as potential population synchronisers and could underly a Moran effect. We considered in our analyses the effect of four weather variables on survival, recruitment, and population growth: cumulative precipitation during breeding (March-May; ‘rainMM’) and non-activity periods (June-February; ‘rainJF’); the minimum (‘tempDFI’) and the maximum temperature (‘tempDFS’) during winter (December-February). The weather variables were collected at meteorological stations (in France, Météo France stations; in Great Britain, UK Meteorological Office) located in the vicinity (from 3 to 25 km) of the surveyed populations. These variables were selected based on the biology of the studied species. During the breeding period, low cumulative rainfall can increase mortality risks of eggs and larvae due to desiccation in amphibians in general (Muths et al. 2017) and in urodeles in particular (Church et al. 2007). On the other hand, high rainfall usually stimulates breeding activities (Wells 2010) that are energetically demanding, which could increase mortality during the reproduction period. During the non-activity period when newts have left the pond and adopted a terrestrial lifestyle, high rainfall and high temperature could increase mortality due to respiratory and energetic constraints (Reading 2007, Griffiths et al. 2010). These effects were investigated in POP1, POP2, POP3 and POP5 (at the whole population level), as well as in two subpopulations of POP1 (POP1.1 and POP1.2) and POP5 (POP5.3 and POP5.4); too few individuals were captured in the other subpopulations to perform analyses. As POP4 has a different phenology due to Mediterranean climatic conditions, we considered the four following variables: the cumulative rainfall during the breeding period (October-April, ‘rainOA’) and the non-activity period (May-September, ‘rainMS’); the minimum (‘tempMSI’) and maximum (‘tempMSS’) monthly temperature during the non-activity period (May-September). Additional information about weather variables (e.g. descriptive statistics, covariation patterns) are provided in Appendix 1.

### Weather and density-dependent effects on survival and recruitment

We examined recapture heterogeneity by performing goodness-of-fit tests in the program U-CARE (Choquet et al. 2009a). First, we performed an overall GOF and detected recapture heterogeneity in only two populations, namely POP1 (*df* = 118, *χ*^2^ = 320.08, *p* < 0.0001) and POP3 (*df* = 19, *χ*^2^ = 35.81, *p* = 0.01; Appendix I). In these two populations, we detected both trap-dependence (POP1, *χ*^2^ = −2.81, *p* = 0.004; POP3, *χ*^2^ = −2.13, *p* = 0.03) and transience (POP1, *χ*^2^ = 4.81, *p* < 0.0001; POP3, *χ*^2^ = 3.46, *p* = 0.0005). We therefore considered recapture heterogeneity in capture-recapture models by including Pledger’s heterogeneity mixtures (Pledger et al. 2003).

To investigate whether survival was influenced by conspecific density and weather variables in each of the five populations, we considered a model with three states *A*_1_, *A*_2_ and *D*, where *A* means alive and *D* dead and where 1 and 2 correspond to the first and second heterogeneity class respectively. We considered two possible field observations, captured or not captured. The model was made of three pieces of information: (1) the vector of initial states probabilities, (2) the survival matrix, and (3) the event matrix linking observations to individual latent states. At their first capture, individuals could be in two states: *A*_1_ and *A*_2_, resulting in the following vector of initial state probabilities:

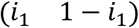

Then, information about survival was updated: individuals in the state *A*_1_ at *t*-1 could reach the state *A*_1_ (i.e. survive) with a probability *ϕ*_1_ or die with a probability 1-*ϕ*_1_; individuals in the state *A*_2_ at *t*-1 could reach the state *A*_2_ with a probability *ϕ*_2_ or die with a probability 1-*ϕ*_2_. This results in the following matrix:

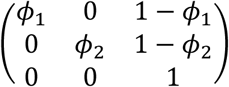

As we are not interested in survival heterogeneity, we hold *ϕ*_1_ and *ϕ*_2_ to be equal in all the models. Lastly, observations were modeled: individuals in the state *A*_1_ could be captured with a probability *p*_1_ or not captured with a probability 1-*p*_1_; individuals in the state *A*_2_ could be captured with a probability *p*_2_ or not captured with a probability 1-*p*_2_. This leads to the following event matrix:

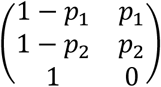

To examine whether annual adult recruitment (i.e., the proportion of sexually mature individuals recruited each year) was affected by conspecific density and weather variables in each of the five populations, we built a model following the structure of Pradel’s (1996) model, in which recruitment was modeled by reversing capture histories and analyzing them backwards. The model is conditional upon the first capture. The recruitment probability Ψ was estimated as the probability that an individual present at time *t* was not present at *t*-1, that is, the proportion of “new” individuals in the population at *t*, while accounting for capture heterogeneity. Note that in our multievent model, survival cannot be estimated along with recruitment. This model has the same structure as the previous model and only the recruitment matrix was modified as following:

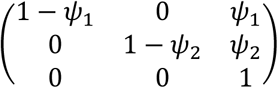

We tested our hypotheses about survival and recruitment separately and by following a similar procedure. The models were implemented in the E-SURGE program (Choquet et al. 2009b). The datasets from the five populations of *T. cristatus* differed in terms of the number of study years and study periods (Appendix S1). Hence, we separately analyzed the five datasets and then compared the population’s responses to weather variations and density using the outputs of the best-fitting models for each population. Models were ranked through a model-selection procedure using Akaike information criterion adjusted for a small sample size (AICc). The analyses were carried out in two steps. First, from the most general model [ϕ(t + sex), p(het + sex)], we evaluated the three following effects: sex (‘sex’), heterogeneity mixture (low vs high capture rates of individuals; ‘het’) and year-specific variation (‘t’). Only additive models were considered in our analyses; interactions were avoided to limit the problem of parameter identifiability and to increase model stability. We also avoided to include year-specific variation on recapture due to recurrent issue of parameter estimates when the model [ϕ(t + sex), p(t + het + sex)] was considered; in many populations, the number of individuals captured each year was relatively low. Yet, time-specific recapture probability was supported in only two populations (POP1 and POP4; Appendix 3). Furthermore, the models [ϕ(t), p(t)] and [ϕ(t), p(.)] provided very similar estimates of survival in POP1 and POP4 (Appendix 3). Therefore, [ϕ(t + sex), p(het + sex)] was kept as general model as time-specific survival is required in ANODEV analyses (see below). We examined all the possible combinations of effect, resulting in the consideration of 16 models per population and subpopulation.

In a second step, after determining the best-fitting model, we examined the effect of weather and density variables on survival probability using ANODEV as recommended in Grosbois et al. (2008). This approach allowed us to evaluate the fit of a model including a single meteorological covariate (*M*_*cov*_) relative that of both the constant (*M*_*cst*_) and the time-dependent (*M*_*t*_) models. The statistic *Ftest*_*cst/cov/t*_ has been derived as following:

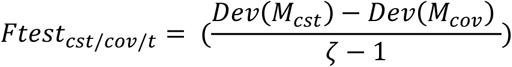

This statistic tests the null hypothesis *H*_*0*_ that the meteorological covariate in *M*_*cov*_ has no significant effect on survival or recruitment. It follows under *H*_*0*_ a Fisher-Snedecor distribution with ζ - 1 and *n* - ζ degrees of freedom. To examine the effect of population density at *t*-1 on survival and recruitment, we estimated population size at *t*-1 by using Horvitz-Thompson estimator. All variables (weather and density) were scaled when they were entered in the models.

### Weather and density-dependent effects on population growth rate

To assess the effect of density-dependence and weather on population growth rate, we used Gompertz state-space (GSS) models (de Valpine & Hastings 2002). We followed the approach described in Kéry & Schaub (2012) and Bancila et al. (2015). The abundance time series of the five populations and the four subpopulations of POP1 and POP5 were analyzed separately. The abundances were corrected by the time-constant recapture probability (i.e. Horvitz-Thompson estimator) obtained from the capture-recapture models. As recapture probability fluctuates over time in several populations (Appendix 3), we verified that using population sizes corrected with time-specific recapture probability did not affect our results (Appendix 5). The log-transformed (log(*X*+1)) population sizes *X*_*t*_ in a pond at time *t* are described by:

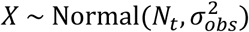

where *N*_*t*_ is the unobserved true population size at time *t* and 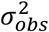 is the observation variance. This piece of the GSS model described the observation process and allows for observation error while assuming that newt abundance may overestimate or underestimate true population size. *N*_*t*_ was defined from a normal distribution:

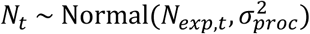

where 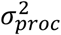 designates the process variance (i.e. stochastic variability in newt abundance) and *N*_*exp,t*_ the expected population size at time *t*. The effect of population density at *t* – 1 was modeled as following:

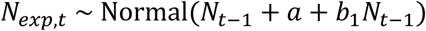

where *a* is the intercept and *b* the coefficient slope for the effect density at *t* – 1. To evaluate the effect of weather variable on population growth rate, we considered models including one weather variable at a time. The model had the following form:

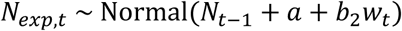

where *w*_*t*_ is the weather covariate at time *t* and *b*_2_ is the slope coefficient. Climatic variables were standardized to a mean 0 and variance 1.

The models were fitted in JAGS (Plummer 2003), using the R package jagsUI (Kellner 2018), with vague normal priors with a mean 0 and a precision of 0.0001 for *a* and *b* parameters. Vague uniform priors ranged within an interval 0-10 were applied for the standard deviation of 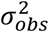 and 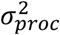. We used a normal prior with a mean equal to the log-transformed population size and a variance of 100 for the first value of the time series. Three MCMC chains were ran with 2 000,000 iterations and a burn-in of 1,000.000. Chains were thinned by a factor 20. We assessed model convergence with the Gelman-Rubin statistic R-hat; we assumed that the model convergence was satisfactory when R-hat values were less than 1.1 (Brooks & Gelman 1998, Gelman & Hill 2006). We considered that the effect of a variable on population growth rate was significant if the 95% CI of the slope coefficient did not include 0.

### Population synchrony within and between-populations

The temporal synchrony among populations was assessed using the variance partitioning methodology (Grosbois et al. 2009, Schaub et al. 2015). More specifically, we built a generalized mixed model (GLMM) treating the estimate of population size as the dependent variable using a Poisson distribution to specify the error term and the ln transformation as the link function. The study population was introduced as a fixed explanatory term in the model. We estimated yearly abundance by dividing the number of captured newts by the time-constant recapture probability obtained from the multievent capture-recapture models described below (i.e. Horvitz-Thompson estimator). For subpopulations for which capture-recapture were not built, we used the mean recapture probability estimated for the whole population. As recapture probability may vary over time in several populations (Appendix 3), we verified that using population sizes corrected with time-specific recapture probability did not affect our results. Given the uncertainty of population size estimates, we also controlled for our estimates of population synchrony not being substantially underestimated. Neither of these adjustments substantially changed our results, and are presented in Appendix 2 for details. Both the time *t* and its interactive effect with the population *t*\**p* were introduced as random effects in the model. The model was thus specified as follows:

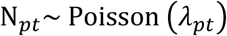

where *N*_*pt*_ is the estimated demographic size of population *p* at time *t* following a Poisson distribution with *λ*_*pt*_, the expected demographic size as

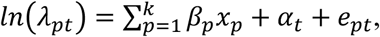

with

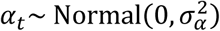

where 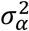 is the shared temporal variance between the *k* populations, and

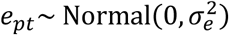

where 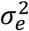 the residual temporal variance specific to each *p* population. *β*_*p*_ is the main population effect (associated to the dummy variable *x*_*p*_ specifying the partial intercept for the population *p*) accounting for the overall difference of demographic size between the *k* populations.

The temporal synchrony among populations was then measured using Intraclass Correlation Coefficient (ICC) that expresses the ratio of the temporal variation that is common across all studied populations and the entire amount (i.e. spatiotemporal) of variation. ICC values close to 1 indicate a quasi-perfect temporal synchrony among populations, while at the opposite, ICC values close to 0 indicate complete asynchrony, and ICC below 0.5 a relatively low synchrony. We computed the adjusted Intraclass Correlation Coefficient (ICC_adj_) using the delta method to accommodate the latent variance inherent to the Poisson distribution associated to the GLM model (see Nakagawa et al. 2017, for justification and implementation):

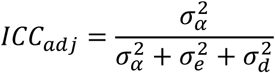

where 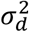 is the latent variance inherent to the Poisson distribution estimated as

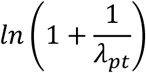

with

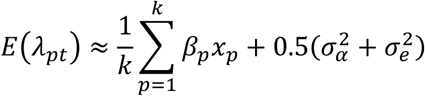

To estimate population synchrony at the continental scale we restricted the time series of each population to the 2000-2015 period in order to ensure a balanced dataset across populations. In the case of POP1 and POP5, we used the estimated total population size to compute this estimation. Estimation of population synchrony at the local scale was performed separately for POP1 and POP5 using the population size estimated for each of their subpopulations (i.e. replacing population by subpopulation in the formula above) during the whole study period.

## RESULTS

### Population synchrony at local and continental scales

Population size drastically varied over time and across populations and subpopulations (**Fig.1**). The level of synchrony at the local scale varied greatly between populations (Table 1): it was relatively high in POP5 but rather low in POP1 as indicated by their respective ICC. At the continental scale, we found a relatively low level of population synchrony as indicated by the estimated ICC (0.212, Table 1). We further used another GLMM to decompose the total variance as the sum of the shared temporal variance between POP1 and POP5, the shared temporal variance between sub-populations of each population and the residual temporal variance specific to each subpopulation. The level of synchrony between populations was estimated at the lower bound (i.e. 0, Appendix 2), and the average level of synchrony between subpopulations (within each population) was moderate (i.e. 0.488, Appendix 2).

**Fig. 1.**
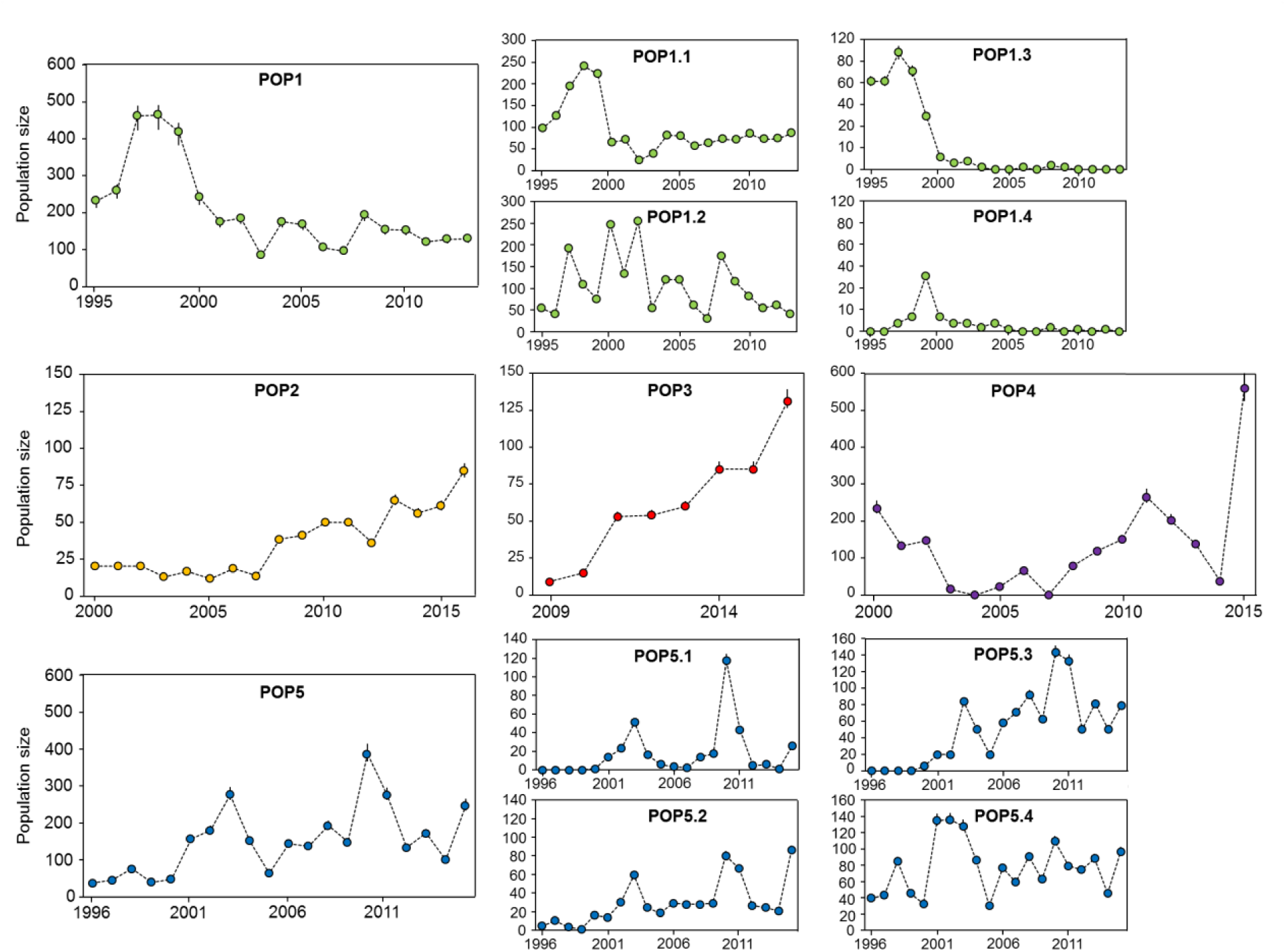
Population sizes and 95% CI (derived from survival models) in the five studied populations (POP1, POP2, POP3 and POP4) of crested newt and their respective subpopulations (POP1: POP1.1, POP1.2, POP1.3 and POP1.4; POP5: POP5.1, POP5.2, POP5.3 and POP5.4). The confidence intervals were obtained by dividing the counts by the upper and lower confidence limits of recapture probability.

**Table 1.**
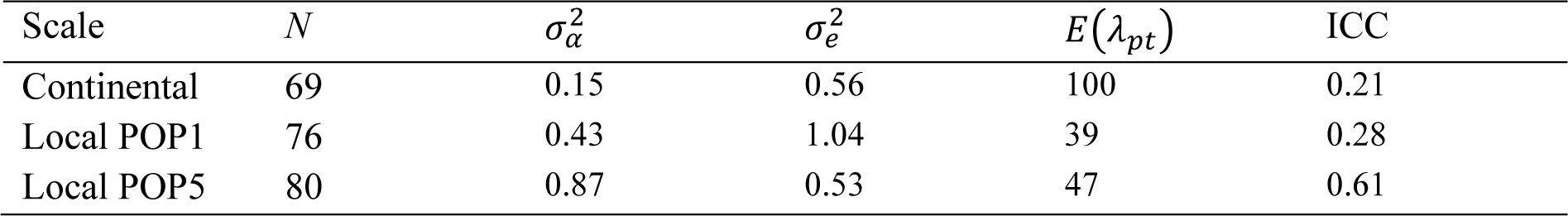
Synchrony among the five populations (i.e. continental level) and within two populations (i.e. local level, POP1 and POP5) of crested newt. ICC and shared/specific temporal variance of population size. N is the total number of observations used, 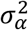 is the shared temporal variance among populations, 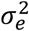 is the temporal variance specific to each population, E(λ) the expected mean demographic size over all populations.

### Survival and recruitment variation within- and between-populations

Within-populations, our analyses revealed that mean survival (provided by the model where survival was constrained to be constant) displayed slight subpopulation-specific variation in POP1 and POP5. In POP1, the survival was 0.61±0.02 in POP1.1 and 0.66±0.02 in POP1.2. In POP5, survival ranged from 0.23±0.01 in POP5.3 to 0.36±0.01 in POP5.4. In the different subpopulations of POP1 and POP5, survival varied widely between years (**Fig.2**). At the population level, survival displayed broad population-specific variation. Mean survival was relatively high in POP2 and POP3, 0.83±0.02 and 0.87±0.02 respectively. Survival was lower (0.62±0.01) in both POP1 and POP4 and drastically lower in POP5 (0.29±0.01). Survival varied between years to a lesser extent in POP2 and POP3 (**Fig.2**) where mean survival was higher. By contrast, it broadly varied between years in POP1, POP4 and POP5 (**Fig.2**). The effect of sex on survival was absent or marginal in all the populations and subpopulations (**Fig.2**, and model selection procedures in Appendix 3).

**Fig. 2.**
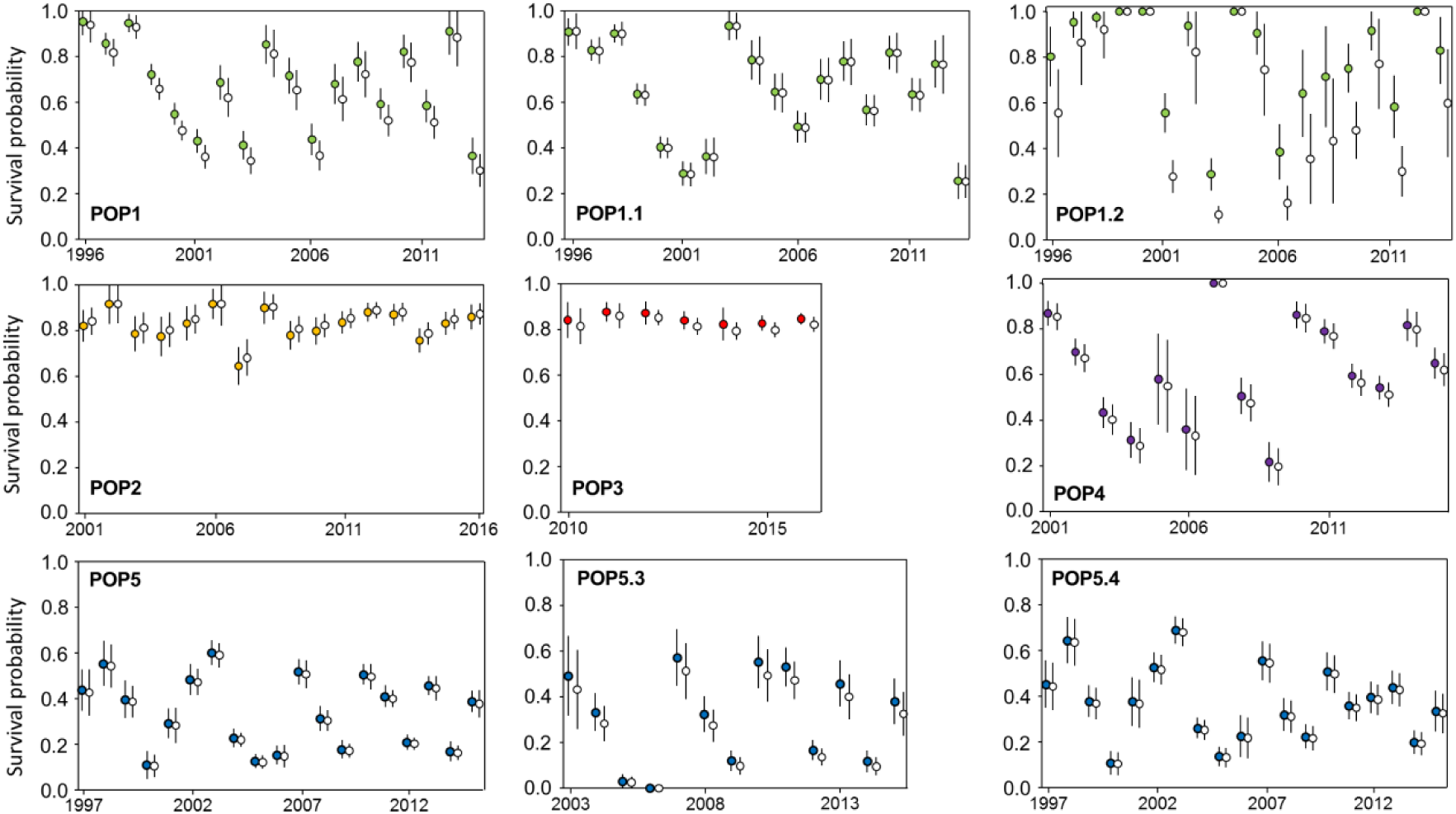
Survival probability in five populations (POP1, POP2, POP3, POP4 and POP5) of crested newt in Western Europe. Survival has been also estimated in subpopulations (i.e. ponds or groups of ponds) of POP1 (POP1.1 and POP1.2) and of POP5 (POP5.3 and 5.4). In subpopulation POP.5.3, we excluded the first five years of survey (1996-2002) because few newts were captured. Males are shown in full circles; females are shown in empty circles.

At the within-population level, recruitment displayed slight subpopulation-specific variation in POP1 and POP5. In POP1, recruitment was 0.35±0.02 in POP1.1 and 0.32±0.02 in POP1.2. In POP5, recruitment was 0.51±0.03 in POP5.3 and 0.65±0.02 in POP5.4. In the different subpopulations of POP1 and POP5, recruitment fluctuated among years (**Fig.3**). At the population level, mean recruitment (estimated in the model where recruitment was constrained to be constant) was relatively similar in POP1 (0.33±0.01), in POP2 (0.25±0.03) and in POP3 (0.29±0.03). Yet, it was higher in POP4 (0.43±0.01) and drastically increased in POP5 (0.71±0.01). In all five populations, recruitment was highly variable between years (**Fig.3**). The sex had a little effect on recruitment in all the populations and subpopulations (**Fig.3**, and model selection procedures in Appendix 4).

**Fig. 3.**
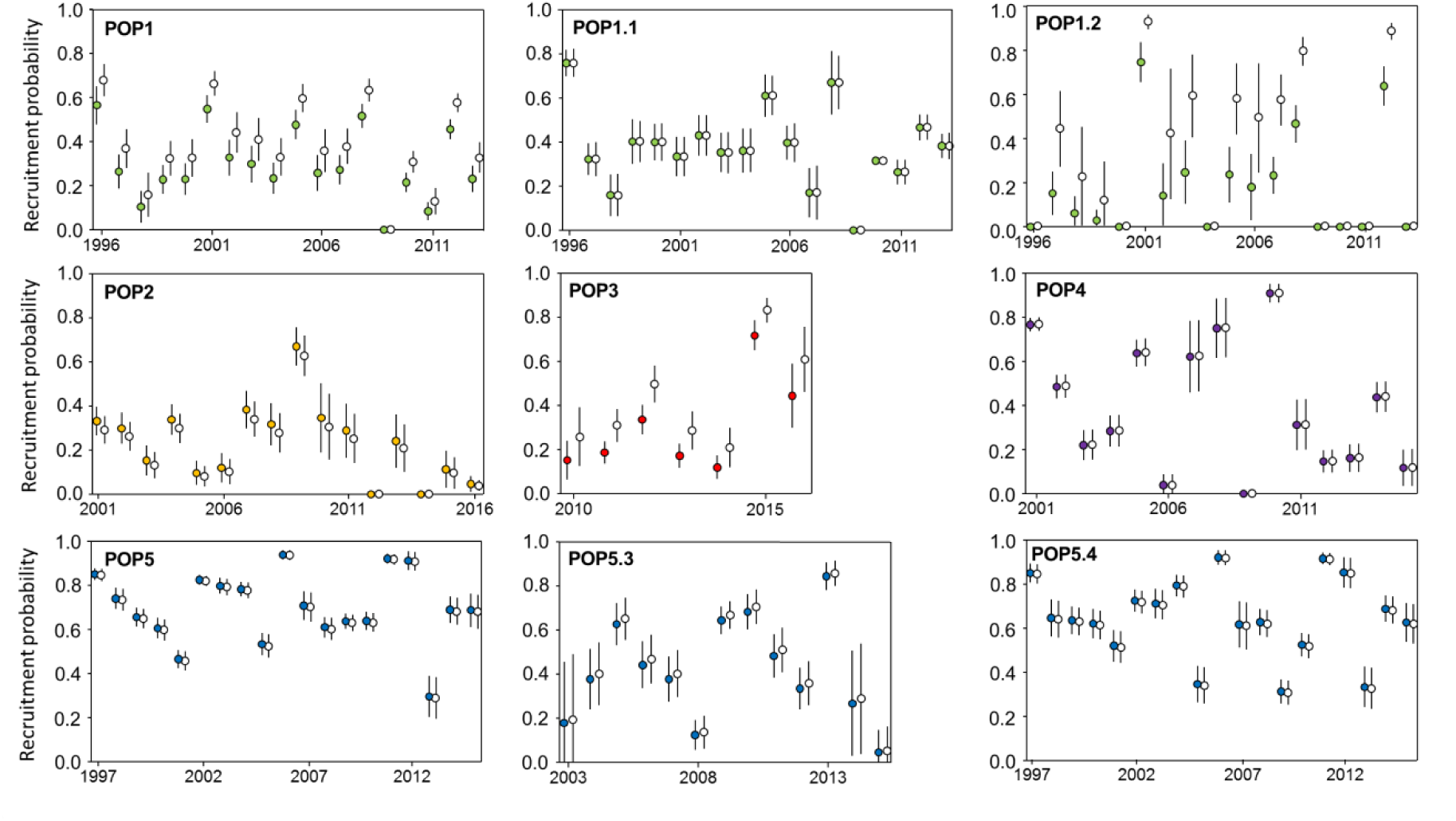
Recruitment probability in five populations (POP1, POP2, POP3, POP4 and POP5) of crested newt in Western Europe. Survival has been also estimated in subpopulations (i.e. ponds or groups of ponds) of POP1 (POP1.1 and POP1.2) and of POP5 (POP5.3 and 5.4). In subpopulation POP.5.3, we excluded the first five years of survey (1996-2002) because few newts were captured. Males are shown in full circles; females are shown in empty circles.

### Influence of density-dependence and weather on survival and recruitment

Our analyses revealed a weak effect of density-dependence and heterogenous effect of weather on survival at both subpopulation and population levels (**Fig.4**) – ANODEV results are summarized in **Fig.5**, and complete ANODEV outputs can be found in Appendix 3. The only significant effects were found in POP1 (**Fig.4**). At the subpopulation level, survival was negatively influenced by the cumulative rainfall during the March-May period (RainMM, *F* = 11.68, *p* = 0.003) and during the June-February period in subpopulation POP1.1 (RainJF, *F* = 5.14, *p* = 0.04) but not in POP1.2. In this subpopulation, we detected a negative effect of population density on survival (density, *F* = 4.73, *p* = 0.04). At the population level, the two weather effects detected in POP1.1 were also detected at whole population level (RainMM, *F* = 12.97, *p* = 0.002; RainJF, *F* = 5.64, *p* = 0.03) but no effect of population density was found. In POP4, we also detected a negative effect of the cumulative rainfall during the non-activity period (RainMM, *F* = 5.91, *p* = 0.03) on survival.

**Fig. 4.**
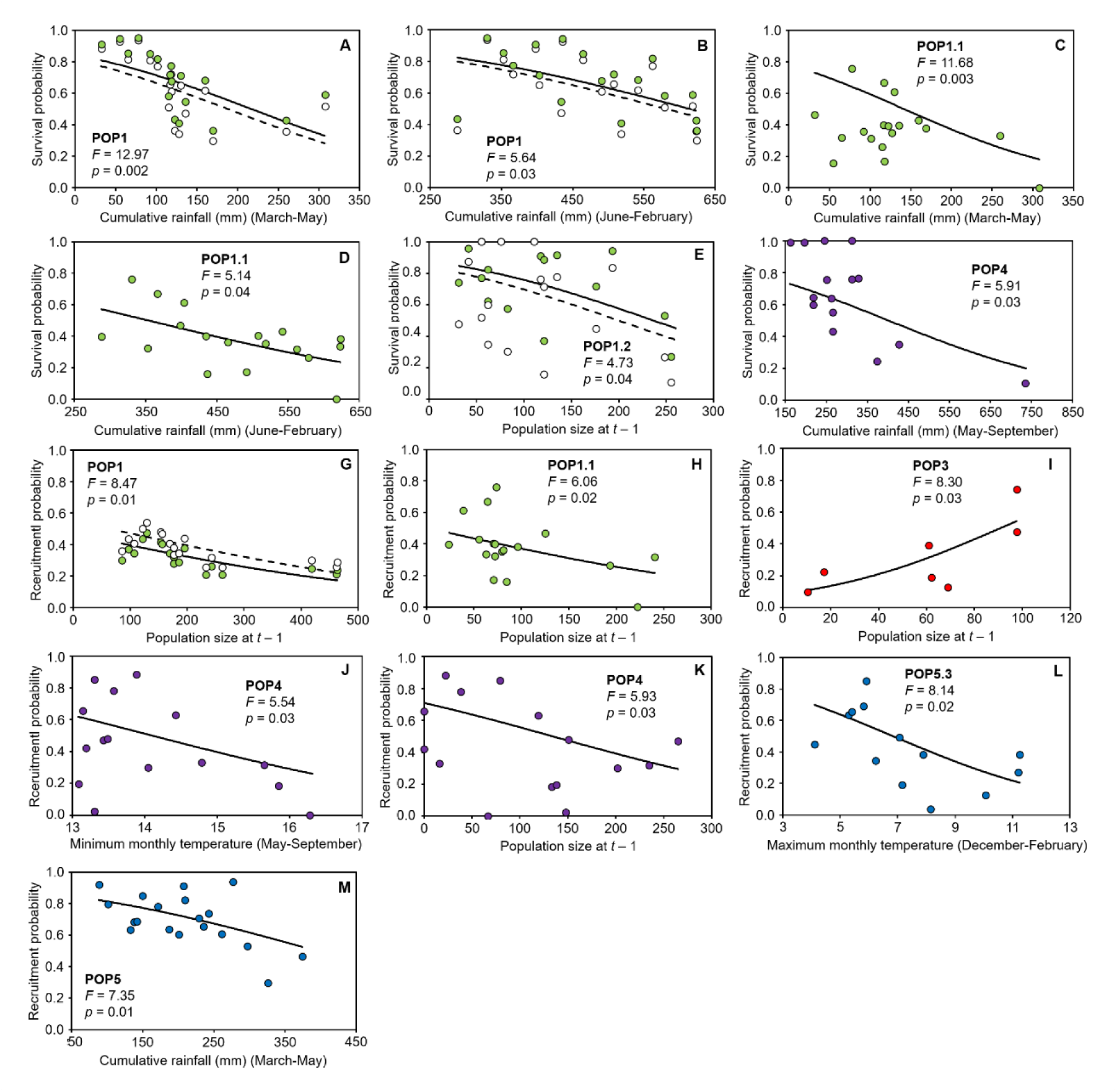
Effects of density (i.e. population size at t–1) and weather variables on survival (A-E) and recruitment (F-J) in five populations of Triturus cristatus in Europe. In the POP1, POP2, POP3 and POP5, four meteorological factors were considered in the analyses: cumulative rainfall during the breeding period (March-May) and the non-activity period (June-February) as well as minimum and maximum monthly temperature during the winter (December-February). As POP4 displays a phenological shift, we considered the four meteorological factors: cumulative rainfall during the breeding period (October-April) and the non-activity period (May-September) as well as minimum and maximum monthly temperature during the non-activity period. The figure shows all the significant relationships (assessed using ANODEVs) between demographic parameters and weather variables. For each relationship, the time-specific estimates of the demographic parameters are fitted against the considered covariate (circles) and lines figure out the predictions provided by the model including the weather covariate. In the cases where a sex-specific effect was retained, parameter estimates and predictions for males are shown in full circles and full lines; those of females are shown in empty circles and broken lines.

**Fig. 5.**
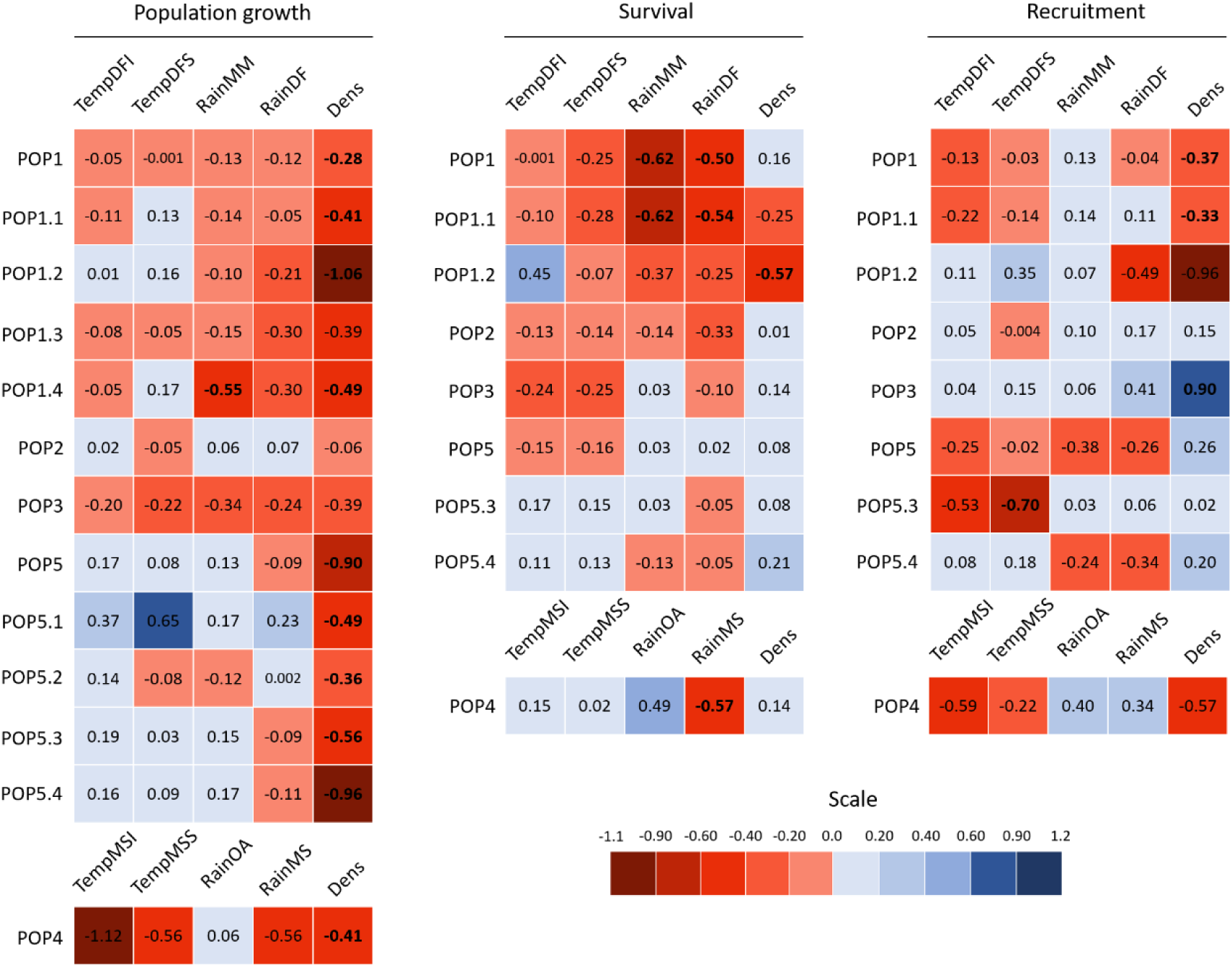
Effects of density (i.e. population size at t–1) and weather variables on population growth survival, and recruitment in five populations of Triturus cristatus (POP1 to POP5) in Europe. POP1 and POP5 includes four subpopulations (POP1.1 to POP1.4, and POP5.1 to POP5.4). In the POP1, POP2, POP3 and POP5, four meteorological factors were considered in the analyses: cumulative rainfall during the breeding period (RainMM) and the non-activity period (RainJF) as well as minimum (TempDFI) and maximum (TempDFS) monthly temperature during the winter. As POP4 displays a phenological shift, we considered the four meteorological factors: cumulative rainfall during the breeding period (RainOA) and the non-activity period (RainMS) as well as minimum (TempMSI) and maximum (TempMSS) monthly temperature during the non-activity period. The coefficient slope of the relationship between demographic rates and variables is given and appear in bold when the effect is significant (capture-models: p-value of the ANODEV < 0.05; GSS models: the 95% CI does not include 0). For recruitment and survival, a complete description of the coefficient slopes (and their 95% CI) and the outputs of ANODEVs is provided in Appendix 3 and 4. The complete outputs of GSS models (posterior distribution, coefficient slopes) are provided in Appendix 5.

Our result revealed heterogeneous effects of weather and density-dependence on adult recruitment (**Fig.4**) – ANODEV results are summarized in **Fig.5**, and complete ANODEV outputs can be found in Appendix 4. At the subpopulation level, we detected a negative density-dependent effect on recruitment in POP1.1 (density, *F* = 6.06, *p* = 0.02) while no effect was detected in POP1.2. In POP5.3, we found a negative relationship between recruitment and the maximum mean temperature during winter (TempDFS, *F* = 8.18, *p* = 0.01) whereas no effect was detected in POP5.4. At the population level, we detected a negative density-dependent effect on recruitment in POP1 (density, *F* = 8.64, *p* = 0.01) and POP4 (density, *F* = 5.93, *p* = 0.03). By contrast, we found a positive effect of density on recruitment in POP3 (density, *F* = 8.30, *p* = 0.03). We detected an effect of weather on recruitment in two populations. In POP5, recruitment was negatively affected by cumulative rainfall during the aquatic period (RainMM, *F* = 7.35, *p* = 0.01); in POP4, the minimum temperature during the non-activity period had a negative impact on recruitment.

### Influence of density-dependence and weather on population growth rate

Negative density-dependence was a common phenomenon within populations (for the GSS outputs, see Appendix 5). In POP5, we found significant negative density-dependent effects on population growth rate in the four subpopulations POP5.1 (*b*_1_ = −0.49±0.29), POP5.2 (*b*_1_ = −0.36±0.18), POP5.3 (*b*_1_ = −0.56±0.22) and POP5.4 (*b*_1_ = −0.96±0.39). In POP1, we detected a negative density-dependent effect on population growth in POP1.2 (*b*_1_ = −1.06±0.43) and a trend (i.e. large size effect, but 95% CI including 0) in subpopulation POP1.1 (*b*_1_ = −0.41±0.29), POP1.3 (*b*_1_ = −0.12±0.08) and POP1.4 (*b*_1_ = −0.39±0.28). By contrast, weather effects were slight and less frequent (**Fig.5**). In POP1, our results indicate that cumulative rainfall during the aquatic period negatively affects population growth rate in subpopulation POP1.4 (*b*_2_ = −0.55±0.23). In addition, we detected a trend of a detrimental effect of cumulative rainfall during the non-aquatic period in POP1.2 (*b*_2_ = −0.21±0.18) and POP1.3 (*b*_2_ = −0.30±0.17); the sign was also negative in POP1.1 (*b*_2_ = −0.24±0.23) and POP1.4 (*b*_2_ = −0.30±0.26). In POP5, we only detected trends of weather effects in the subpopulation POP5.1 where population growth was positively affected by the minimum temperature during winter (*b*_2_ = 0.16±0.12) and by rainfall during the activity period (*b*_2_ = 0.17±0.14).

At the population level, density-dependent regulation was also widespread (for the GSS outputs, see Appendix 5). We detected significant negative effects of population density on population growth in POP4 (*b*_1_ = −0.91±0.46) and POP5 (*b*_1_ = −0.41±0.23). In addition, a similar trend was found in POP1 (*b*_1_ = −0.28±0.24) and POP3 (*b*_1_ = −0.39±0.30). However, we failed to detect any effect of density-dependence in POP2 (*b*_1_ = −0.06±0.18). By contrast, weather effects on population growth were slighter and scarcer (**Fig.5**). In POP1, we detected a trend of a negative relationship between population growth and the cumulative rainfall during both the aquatic period (*b*_2_ = −0.13±0.10) and the non-aquatic period (*b*_2_ = −0.12±0.09). A similar trend was also found in POP3 where population growth rate was negatively affected by the cumulative rainfall during the aquatic period (*b*_2_ = −0.34±0.25). In POP4, we found a trend of a negative relationship between population growth and the minimum temperature during the non-aquatic (aestivation) period (*b*_2_ = −1.11±0.63).

## DISCUSSION

We found a low level of demographic synchrony at both local and continental levels. Our study also revealed that weather had weak but spatially variable effects on survival, recruitment and population growth rate. In contrast, density-dependence was a common phenomenon (at least for population growth rate) in all the populations and subpopulations. Taken together, our results suggest that population dynamics more closely depend on habitat-specific factors rather than regional weather fluctuations, resulting in a low synchrony within and between populations.

### Weak and spatially heterogeneous effects of weather on population dynamics

Overall, we found that weather variation has relatively weak effects on demographic rates at the subpopulation and populations levels. At the population scale, we detected only three significant relationships between survival and weather factors (cumulative rainfall during the breeding and non-activity periods) in two populations (POP1 and POP4). In POP1, the two relationships (rainfall during the breeding and non-aquatic periods) were also detected in one of the two subpopulations (POP1.1). The negative relationship between survival and rainfall during non-breeding activity (POP1 and POP4) confirms the outcomes of a previous study over a shorter timeframe on this population (Griffiths et al. 2010). High rainfall often results in waterlogged soils, which could impair respiration during overwintering. It could also stimulate newt activity during periods with unsuitable weather conditions (i.e. summer in POP4), resulting in a negative impact on their survival. Moreover, our results indicate that high rainfall during the breeding period also negatively affects adult survival (POP1). As in many other amphibians, high rainfall levels during the reproduction period increases the level of energetically-demanding activities related to breeding (breeding migration, mate searching activities, competition for mates; Wells 2010), which may increase mortality rates. Our findings therefore suggest that these weather effects on adult survival were highly context-dependent and broadly inconsistent across the distribution range of *T. cristatus*. This confirms recent work that highlighted strong context-dependent effects of weather on survival in other amphibian species (Cayuela et al. 2016a, 2017; Muths et al. 2017).

Our study also revealed that weather had a low influence on adult recruitment. Indeed, we detected only two significant relationships between adult recruitment and one weather factor in POP5 and POP4. In POP5, we found a negative relationship between recruitment and cumulative rainfall during the breeding period at year y-1. A higher level of energetically-demanding activities in immatures, such as natal dispersal during the previous year, might increase mortality rates. Such increases could then result in lower recruitment the following year. We also recorded a negative relationship between recruitment and the maximum monthly temperature during the non-activity period in POP4 and the subpopulation POP5.3. This pattern is in accordance with a previous study in amphibians showing that higher temperatures may increase mortality rates due the depletion of energy reserves during overwintering (e.g. Reading 2007). Lower survival of immatures at year y-1 could then affect recruitment. Like survival, the weather effects on adult recruitment varied across the distribution range of *T. cristatus*.

Finally, GSS models did not reveal any significant effect (only trends) of weather on population growth rate at either subpopulation or population levels. This suggests that the three weather variables considered in our study that have been previously reported as important drivers for amphibians (e.g. Cayuela et al. 2016a, Muths et al. 2017), have a relatively weak effect on *T. cristatus* population dynamics. It is also congruent with our capture-recapture analyses that showed slight and spatially heterogeneous effects of weather on adult survival and recruitment. The spatial inconsistency of the weather effects within and between-population levels is thus a potential mechanism that might reduce the population synchrony at landscape and continental scales.

### Density-dependence, a central driver of population dynamics

Our study revealed that density-dependence during the adult stage was a common driver in the dynamics of *T. cristatus* populations. We detected negative effects of density on population growth rates in almost all the subpopulations and the populations, although the detection of density-dependent effects on survival and recruitment was more variable. We only found a negative effect of density on survival in subpopulation POP1.2. Yet, we detected a negative effect of density on adult recruitment in two of the five studied populations (POP1 and POP4). By contrast, we detected a positive effect on density dependence on recruitment in POP3. This population occupies a pond created at the beginning of the survey, and positive density dependence likely results from mate finding Allee (Gascoigne et al. 2009, Cayuela et al. 2019). These results suggest that the strength of the density-dependent effects could be higher before than after metamorphosis, confirming the conclusions of experimental studies (Altwegg 2003, but see Patrick et al. 2008). Although individual data on subadults were not available in our study, the negative effects of conspecific density on survival and growth at both larval and juvenile stages is common in amphibians (e.g. Altwegg & Reyer 2003, Harper & Semlitsch 2007). Intraspecific competition at larval and juvenile stages is therefore probably involved in the widespread negative effects of density on population growth rate detected in the five studied populations of *T. cristatus*. As no dispersal events occur among populations (and are rare in adults within populations), we therefore assume that populations, as well as subpopulation in POP1 and POP5, should have independent density-dependent structure and that density-dependence is a potential mechanism of population asynchrony at landscape and continental scales.

### Weak population synchrony within and between-populations

A higher level of population synchrony was observed within-populations than between-populations. However, population synchrony was relatively weak, excluding a strong Moran’s effect, even at a relatively small spatial scale: the correlation coefficients were relatively low among the subpopulations of POP1 (0.28) and POP5 (0.61). A higher ICC in POP5 than in POP1 is likely due to lower Euclidean distances between subpopulations (from 200 and 800 m in POP1 and from 60 to 430 m in POP5). At this spatial scale, the meteorological conditions and the landscape characteristics are expected to be relatively homogeneous. Therefore, the low subpopulation synchrony necessarily results from the effects of local factors such as asymmetric density-dependent structures between ponds (e.g. differences in the carrying capacities of ponds) and local environmental variation (e.g. low habitat quality and anthropogenic disturbance). Our results showed that density-dependence is a common phenomenon in subpopulations of POP1 and POP5, which likely reduce their synchrony level in the absence of high dispersal flows. In addition, pond-specific variation in survival, recruitment and subpopulation size (Unglaub et al. 2015, 2018) also probably reduces the level of subpopulation synchrony. At the continental scale, the level of population synchrony was particularly weak (i.e. 0.2 when performing the analysis only at the continental scale, close to 0 when performing the analysis at both the continental and local scale using POP1 and POP5, see appendix 2). In addition to the mechanisms discussed above, regional climatic variation could further reduce population synchrony. Moreover, the very low level of synchrony could be also caused by inter-population differences in demographic strategies (in amphibians, see for instance Cayuela et al. 2016b). Indeed, our results show broad variation in survival and recruitment rates, several presenting ‘fast’ life histories (low survival and high recruitment, POP1) while other displaying decelerated life histories (high survival and low recruitment, POP2 and POP3).

### Conclusion

Our study revealed that weather effects were weak and highly variable within and between populations. By contrast, density-dependence was a central mechanism in *T. cristatus* population dynamics. Our study indicates that these two demographic mechanisms are probably involved in the low population synchrony observed within populations and between populations at continental level. To date, our study is one of the few that have linked population synchrony, density-dependence and heterogeneous demographic responses to weather in natural populations. Although they are focused on an amphibians, our findings support the idea that Moran’s effect is low in species (e.g. many freshwater insect and fishes; see for instance Chevalier et al. 2015) where the population dynamics more closely depends on local factors (e.g. population density and habitat characteristics) than on large-scale environmental fluctuation (e.g. regional climatic variation). A relatively low level of synchrony likely has far-reaching consequences for the long-term viability of the spatially structured populations of these species. Especially, low population synchrony likely limits the risk of extinction of the spatially structured populations, allowing rescue effects between subpopulations and the colonization of patches after local extinction. It probably has important consequences for population response to climate change, by mitigating extinction risk caused by large-scale climatic anomalies. To generalize our findings, we encourage further studies to examine the relative contribution of synchronizing (dispersal, weather variation) and desynchronizing (density, local environmental factors) on population synchrony and dynamics in broader range of taxa.

## Acknowledgements

We thank the many staff, students and volunteers who have contributed to data collection in populations from England over the past 25 years, particularly C. Williams, S. Young, D. Sewell, B. Lewis, A. Wright, P. Walsh, J. Phillips. All surveys in England were carried out under licence from Natural England. We also warmly thank the Conservatoire d’Espaces Naturels Languedoc-Roussillon who collected the data in POP5, especially Thomas Gendre and Pauline Bernard.

RAG, NZ, JWA, PP and PJ collected the data; HC and JPL analysed the data; HC and AB led the writing of the manuscript. All authors contributed critically to the drafts and gave final approval for publication.

