## Appendix S1 for "Drivers of amphibian population dynamics and asynchrony at local and continental scales"

**Appendix 1: Capture-recapture data and environmental variables**

**Fig. S1.1**. Map of the study area showing the five populations of *Triturus cristatus*.

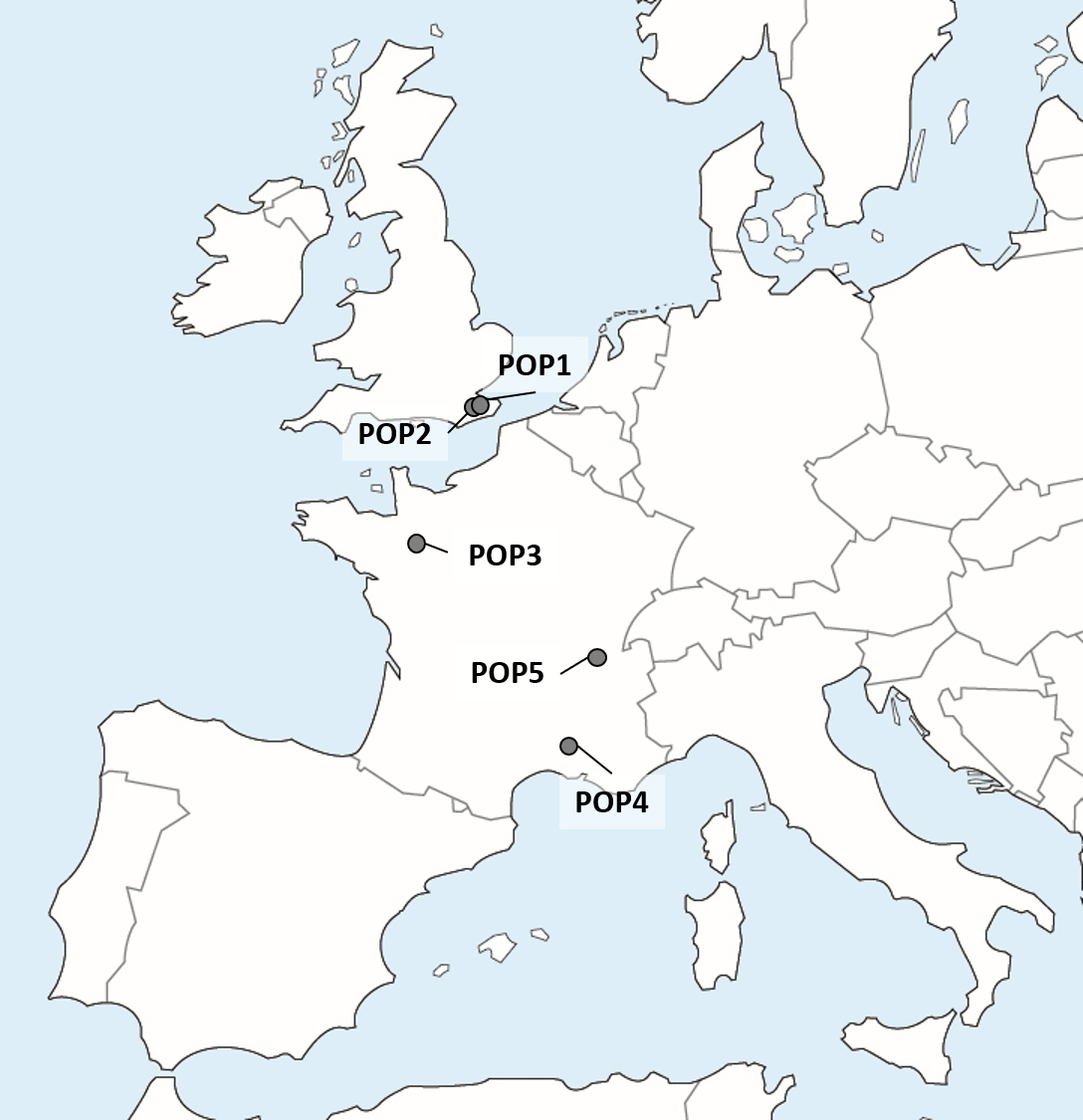

**Table S1.1**. Survey duration and period in the five populations of *T. cristatus*.

| Sex | POP1 | POP2 | POP3 | POP4 | POP5 |
| --- | --- | --- | --- | --- | --- |
| Survey duration | 19 years | 17 years | 8 years | 16 years | 20 years |
| Period | 1995-2013 | 2000-2016 | 2009-2016 | 2000-2015 | 1996-2015 |

**Table S1.2**. Number of females and males identified during the capture-recapture survey in the five populations of *T. cristatus*.

| Sex | POP1 | POP2 | POP3 | POP4 | POP5 |
| --- | --- | --- | --- | --- | --- |
| Female | 577 | 54 | 134 | 465 | 1096 |
| Male | 653 | 72 | 80 | 391 | 1186 |

**Table S1.3**. Number of females and males identified during the capture-recapture survey in the four subpopulations of POP1 and POP5.

| Sex | POP1.1 | POP1.2 | POP1.3 | POP1.4 | POP5.1 | POP5.2 | POP5.3 | POP5.4 |
| --- | --- | --- | --- | --- | --- | --- | --- | --- |
| Female | 289 | 183 | 58 | 27 | 117 | 212 | 414 | 353 |
| Male | 382 | 190 | 55 | 26 | 134 | 200 | 511 | 341 |

**Table S1.4**. Goodness-of-fit (GOF) test. Overall GOF test for recapture heterogeneity. The significant tests are shown in bold.

| Populations | Global test |
| --- | --- |
| POP1 | ***df* = 118,** $\boldsymbol{\chi}^{\boldsymbol{2}}$ **= 320.08, p < 0.0001** |
| POP2 | *df* = 52, $\chi^{2}$ = 22.51, p = 0.99 |
| POP3 | ***df* = 19,** $\boldsymbol{\chi}^{\boldsymbol{2}}$ **= 35.81, p = 0.01** |
| POP4 | *df* = 56, $\chi^{2}$ = 56.89, p = 0.44 |
| POP5 | *df* = 55, $\chi^{2}$ = 46.76, p = 0.77 |

**Table S1.5**. Goodness-of-fit (GOF) test. GOF test for transience and trap-dependence. The significant tests are shown in bold.

| Populations | Transience | Trap-dependence |
| --- | --- | --- |
| POP1 | $\boldsymbol{\chi}^{\boldsymbol{2}}$ **= 4.08, p < 0.0001** | $\boldsymbol{\chi}^{\boldsymbol{2}}$ **= -2.81, p = 0.004** |
| POP3 | $\boldsymbol{\chi}^{\boldsymbol{2}}$ **= 3.46, p = 0.0005** | $\boldsymbol{\chi}^{\boldsymbol{2}}$ **= -2.13, p = 0.03** |

**Table S1.6**. Mean and standard deviation of weather variables.

| Variable | POP1 | POP2 | POP3 | POP5 |
| --- | --- | --- | --- | --- |
| TempDFS | 7.6 (1.2) | 7.9 (1.3) | 2.1 (1.3) | 0.1 (1.2) |
| TempDFI | 2.6 (1.0) | 2.9 (1.1) | 8.6 (1.3) | 7.4 (2.3) |
| RainMM | 127.6 (66.9) | 135.4 (66.6) | 177.7 (78.3) | 208.5 (76.5) |
| RainJF | 474.2 (105.3) | 512.4 (106.2) | 725.4 (113.1) | 691.8 (95.2) |

| Variable | POP4 |
| --- | --- |
| tempMSS | 27.8 (1.1) |
| tempMSI | 14.1 (1.1) |
| RainMS | 564.6 (150.2) |
| RainOA | 304.8 (137.1) |

**Table S1.7**. Weather variables in POP1, correlation matrix.

|  | TempDFS | TempDFI | RainMM | RainJF |
| --- | --- | --- | --- | --- |
| TempDFS | 1.0000000 | 0.90581992 | -0.16322654 | -0.10189107 |
| TempDFI | 0.9058199 | 1.00000000 | -0.05414003 | -0.01719627 |
| RainMM | -0.1632265 | -0.05414003 | 1.00000000 | 0.64067469 |
| RainJF | -0.1018911 | -0.01719627 | 0.64067469 | 1.00000000 |

**Table S1.8**. Weather variables in POP2, correlation matrix.

|  | TempDFS | TempDFI | RainMM | RainJF |
| --- | --- | --- | --- | --- |
| TempDFS | 1.00000000 | 0.92307763 | -0.3277393 | -0.06787043 |
| TempDFI | 0.92307763 | 1.00000000 | -0.2905367 | -0.08056094 |
| RainMM | -0.32773929 | -0.29053674 | 1.0000000 | 0.47407167 |
| RainJF | -0.06787043 | -0.08056094 | 0.4740717 | 1.00000000 |

**Table S1.9**. Weather variables in POP3, correlation matrix.

|  | TempDFS | TempDFI | RainMM | RainJF |
| --- | --- | --- | --- | --- |
| TempDFS | 1.0000000 | 0.9603728 | 0.5913653 | 0.1230146 |
| TempDFI | 0.9603728 | 1.0000000 | 0.5797095 | 0.1506000 |
| RainMM | 0.5913653 | 0.5797095 | 1.0000000 | 0.1858207 |
| RainJF | 0.1230146 | 0.1506000 | 0.1858207 | 1.0000000 |

**Table S1.10**. Weather variables in POP4, correlation matrix.

|  | TempMSS | TempMSI | RainMS | RainOA |
| --- | --- | --- | --- | --- |
| TempMSS | 1.0000000 | 0.5026357 | -0.3124218 | -0.2725628 |
| TempMSI | 0.5026357 | 1.0000000 | -0.5069901 | -0.6320887 |
| RainMS | -0.3124218 | -0.5069901 | 1.0000000 | 0.3971801 |
| RainOA | -0.2725628 | -0.6320887 | 0.3971801 | 1.0000000 |

**Table S1.11**. Weather variables in POP5, correlation matrix.

|  | TempDFS | TempDFI | RainMM | RainJF |
| --- | --- | --- | --- | --- |
| TempDFS | 1.0000000 | 0.63882874 | 0.11540596 | 0.3087676 |
| TempDFI | 0.6388287 | 1.00000000 | 0.03610936 | 0.1280505 |
| RainMM | 0.1154060 | 0.03610936 | 1.00000000 | -0.3102695 |
| RainJF | 0.3087676 | 0.12805052 | -0.31026952 | 1.0000000 |
