## Appendix S2 for "Drivers of amphibian population dynamics and asynchrony at local and continental scales"

**Appendix 2: Population synchrony analysis**

In this appendix, we provide the synchrony analysis performed at both local and continental scale using POP1 and POP5. Results of this analysis are presented in table 2.1. We also provide supplementary analyses taking into account the uncertainty of the population size estimates. The results of these latter analyses are presented in table 2.2 for those performed separately at the continental scale and the local scale, and in table 2.3 for the one performed conjointly at the continental and local scale using POP1 and POP5. We also provide additional results concerning the analysis at the continental scale using population size estimated from time dependent model, these results are reported in table 2.4.

**Synchrony analysis at both local and continental scale**

In order to assess simultaneously the level of population synchrony at both local (i.e. within population) and continental scale (i.e. between populations), we focused our analysis on the time series of the populations of both POP1 and POP5 for which four subpopulations were monitored. We restricted our analysis to the common study period 1996-2013. In order to estimate the ICC at each scale, we simply modified our parameterization of random effects introduced in the GLMM by adding a third random effect. The structure of the GLMM was thus as follows:

$ln\left( \lambda_{mpt} \right)=\sum_{p=1}^{k} \beta_{p}x_{p}+\alpha_{t}+\gamma_{mt}+e_{pt}$,

with $\alpha_{t}\sim Normal\left( 0,\sigma_{\alpha}^{2} \right)$ where $\sigma_{\alpha}^{2}$ is the shared temporal variance between POP1 and POP5, $\gamma_{mt}\sim Normal\left( 0,\sigma_{\gamma}^{2} \right)$ where $\sigma_{\gamma}^{2}$ is the shared temporal variance between subpopuations of the same population, and $e_{pt}\sim Normal\left( 0,\sigma_{e}^{2} \right)$ where $\sigma_{e}^{2}$ is the specific temporal variance of each subpopulation. Note that p, refer in the formula to the sub-population, and m to the population.

Given the hierarchized structure of the random effects, the level of synchrony at the population level can be measured as ${ICC}_{pop}=\frac{\sigma_{\alpha}^{2}}{\sigma_{\alpha}^{2}+\sigma_{\gamma}^{2}+\sigma_{e}^{2}+\sigma_{d}^{2}}$ and the level of synchrony at the sub-population level can be measured as ${ICC}_{subpop}=\frac{\sigma_{\gamma}^{2}}{\sigma_{\gamma}^{2}+\sigma_{e}^{2}+\sigma_{d}^{2}}$

The variance component estimates, the expected mean population size and the two ICC estimates are presented in the following table. As indicated by the results, the shared temporal variance between populations is estimated at the lower bound resulting in a null synchronization between populations (${ICC}_{pop}=0$). There is however a substantial synchronization among sub-populations of the same population, as indicated by the ${ICC}_{subpop}$ (0.49) which roughly correspond to the mean of the local ICC estimated separately for each population (using the whole time series for each one).

**Table 2.1.** Synchrony analyses performed conjointly at both the continental and local scale using POP1 and POP5. *N* is the total number of observations used, $\sigma_{\alpha}^{2}$ is the shared temporal variance among populations, $\sigma_{\gamma}^{2}$ is the shared temporal variance among subpopulations, $\sigma_{e}^{2}$ is the temporal variance specific to each subpopulation, $E\left( \lambda_{mpt} \right)$ the expected mean demographic size over both two populations, ${ICC}_{pop}$ is the ICC at the population level, and ${ICC}_{subpop}$ is the ICC at the subpopulation level.

| *N* | $\sigma_{\alpha}^{2}$ | $\sigma_{\gamma}^{2}$ | $\sigma_{e}^{2}$ | $E\left( \lambda_{mpt} \right)$ | ${ICC}_{pop}$ | ${ICC}_{subpop}$ |
| --- | --- | --- | --- | --- | --- | --- |
| 144 | 0 | 0.7226 | 0.7317 | 41.58 | 0 | 0.489 |

**Synchrony analysis at the population level taking into account the measurement error of the population size.**

In order to examine the sensitivity of our synchrony analyses to the measurement error of the population size estimates, we first derived the measurement error from the IC95% of the population size estimated by the Horwitz-Thompson method. All measurement errors less than 1 were rounded to 1 in order to permit model convergence. These values were then introduced as known variance parameters in the residual matrix of the GLMM, resulting in equality constraint for each population size. The rest of the models were kept similar to the ones used in the previous analyses.

**Table 2.2** Synchrony analyses performed separately at the continental scale (i.e. on the five populations) and the local scale (i.e. within POP1 and POP5) and taking into account the uncertainty of population size estimates. *N* is the total number of observations used, $\sigma_{\alpha}^{2}$ is the shared temporal variance among populations, $\sigma_{e}^{2}$ is the temporal variance specific to each population, $E\left( \lambda_{pt} \right)$ the expected mean demographic size over all populations.

| *SCALE* | *N* | $\sigma_{\alpha}^{2}$ | $\sigma_{e}^{2}$ | $E\left( \lambda_{pt} \right)$ | ICC |
| --- | --- | --- | --- | --- | --- |
| Continental | 69 | 0.17 | 0.72 | 130 | 0.19 |
| Local POP1 | 76 | 0.46 | 0.97 | 37 | 0.31 |
| Local POP5 | 80 | 0.92 | 0.45 | 43 | 0.66 |

**Table 2.3** Synchrony analyses performed conjointly at both the continental and local scale using POP1 and POP5 and taking into account the uncertainty of population size estimates. See the legend of the Table 2.1. for details.

| *N* |  | $\sigma_{\alpha}^{2}$ | $\sigma_{\gamma}^{2}$ | $\sigma_{e}^{2}$ | $E\left( \lambda_{mpt} \right)$ | ${ICC}_{pop}$ | ${ICC}_{subpop}$ |
| --- | --- | --- | --- | --- | --- | --- | --- |
| 144 |  | 0 | 0.54 | 0.49 | 39 | 0 | 0.51 |

**Synchrony analysis at the continental scale using population sizes estimated from the time dependent CMR model.**

The model used is similar to the one detailed in the statistical section. Uncertainty of the population sizes estimates cannot be derived from the CMR model, and are thus not included in the ICC estimation. As the uncertainty of the estimates should be substantially higher than the one derived from time constant CMR models, the ICC calculated is likely downward biaised.

**Table 2.4** Synchrony analysis performed at the continental scale using population size estimated from the time dependent CMR model.

| *SCALE* | *N* | $\sigma_{\alpha}^{2}$ | $\sigma_{e}^{2}$ | $E\left( \lambda_{pt} \right)$ | ICC |
| --- | --- | --- | --- | --- | --- |
| Continental | 69 | 0.11 | 0.90 | 169 | 0.11 |
