## Appendix S4 for "Drivers of amphibian population dynamics and asynchrony at local and continental scales"

**Appendix 4: Effect of density-dependence and local weather on recruitment**

In this Appendix are provided the model selection procedures and the ANODEVs performed for recruitment analyses.

1. *Examining recruitment at the population level*

In this first part of the appendix, we provide analyses carried out separately for each of the five studied populations of *Triturus cristatus* (POP1, POP2, POP3, POP4 and POP5). First, for each population, we defined the best-supported model without weather covariate using Akaike information criteria adjusted for small sample size (AICc) and relative AICc weights (**Table S3.1**). We tested all the possible combination of effects, resulting in 16 models per populations.

**Table S4.1**. Recruitment and recapture in five populations of *Triturus cristatus* in Western Europe: model selection procedure. α = recruitment, p = recapture, AICc = Akaike information criterion adjusted for small sample size, w = relative AICc weights, k = numbers of parameters.

POP1

| Model | k | Dev | AICc | Delta | w |
| --- | --- | --- | --- | --- | --- |
| α(t + sex), p(het + sex) | 23 | 3787.12 | 3833.66 | 0.00 | 0.99 |
| α(t), p(het + sex) | 22 | 3799.70 | 3844.19 | 10.53 | 0.01 |
| α(t), p(het) | 21 | 3808.92 | 3851.36 | 17.70 | 0.00 |
| α(t + sex), p(het) | 22 | 3807.59 | 3852.08 | 18.42 | 0.00 |
| α(t + sex), p(sex) | 21 | 3886.79 | 3929.23 | 95.57 | 0.00 |
| α(t), p(sex) | 20 | 3893.35 | 3933.76 | 100.10 | 0.00 |
| α(sex), p(het + sex) | 6 | 3931.99 | 3944.03 | 110.38 | 0.00 |
| α(.), p(het + sex) | 5 | 3934.13 | 3944.16 | 110.50 | 0.00 |
| α(.), p(het) | 4 | 3940.03 | 3948.05 | 114.39 | 0.00 |
| α(t), p(.) | 19 | 3910.25 | 3948.62 | 114.96 | 0.00 |
| α(sex), p(het) | 5 | 3939.96 | 3949.99 | 116.33 | 0.00 |
| α(t + sex), p(.) | 20 | 3910.12 | 3950.53 | 116.87 | 0.00 |
| α(.), p(sex) | 3 | 4019.36 | 4025.38 | 191.72 | 0.00 |
| α(sex), p(sex) | 4 | 4018.66 | 4026.67 | 193.02 | 0.00 |
| α(.), p(.) | 2 | 4029.53 | 4033.53 | 199.87 | 0.00 |
| α(sex), p(.) | 3 | 4029.16 | 4035.18 | 201.52 | 0.00 |

POP2

| Model | k | Dev | AICc | Delta | w |
| --- | --- | --- | --- | --- | --- |
| α(t + sex), p(sex) | 19 | 770.51 | 810.10 | 0.00 | 0.39 |
| α(t), p(sex) | 18 | 773.34 | 810.76 | 0.67 | 0.28 |
| α(t + sex), p(het + sex) | 21 | 767.86 | 811.80 | 1.71 | 0.17 |
| α(t), p(het + sex) | 20 | 770.64 | 812.39 | 2.30 | 0.12 |
| α(t + sex), p(het) | 20 | 774.26 | 816.02 | 5.92 | 0.02 |
| α(t), p(het) | 19 | 778.37 | 817.95 | 7.86 | 0.01 |
| α(t + sex), p(.) | 18 | 781.43 | 818.86 | 8.76 | 0.00 |
| α(t), p(.) | 17 | 785.41 | 820.68 | 10.58 | 0.00 |
| α(.), p(het + sex) | 5 | 822.17 | 832.29 | 22.19 | 0.00 |
| α(sex), p(sex) | 4 | 825.12 | 833.20 | 23.10 | 0.00 |
| α(sex), p(het + sex) | 6 | 823.32 | 835.49 | 25.39 | 0.00 |
| α(.), p(sex) | 3 | 829.35 | 835.40 | 25.30 | 0.00 |
| α(sex), p(het) | 5 | 832.34 | 842.46 | 32.36 | 0.00 |
| α(sex), p(.) | 3 | 838.38 | 844.43 | 34.34 | 0.00 |
| α(.), p(het) | 4 | 837.76 | 845.85 | 35.75 | 0.00 |
| α(.), p(.) | 2 | 844.03 | 848.05 | 37.96 | 0.00 |

POP3

| Model | k | Dev | AICc | Delta | w |
| --- | --- | --- | --- | --- | --- |
| α(t), p(het) | 10 | 694.48 | 714.93 | 0.00 | 0.93 |
| α(t), p(het + sex) | 11 | 698.83 | 721.38 | 6.45 | 0.04 |
| α(t + sex), p(het + sex) | 12 | 698.18 | 722.83 | 7.90 | 0.02 |
| α(t + sex), p(het) | 11 | 702.66 | 725.21 | 10.27 | 0.01 |
| α(t + sex), p(.) | 9 | 707.05 | 725.42 | 10.49 | 0.00 |
| α(t + sex), p(sex) | 10 | 707.05 | 727.50 | 12.57 | 0.00 |
| α(t), p(.) | 8 | 714.18 | 730.47 | 15.54 | 0.00 |
| α(t), p(sex) | 9 | 714.04 | 732.41 | 17.48 | 0.00 |
| α(sex), p(het + sex) | 6 | 739.79 | 751.97 | 37.03 | 0.00 |
| α(sex), p(het) | 5 | 741.82 | 751.95 | 37.01 | 0.00 |
| α(.), p(het) | 4 | 746.64 | 754.72 | 39.79 | 0.00 |
| α(.), p(het + sex) | 5 | 745.32 | 755.45 | 40.51 | 0.00 |
| α(sex), p(.) | 3 | 756.50 | 762.55 | 47.62 | 0.00 |
| α(sex), p(sex) | 4 | 756.50 | 764.58 | 49.64 | 0.00 |
| α(.), p(.) | 2 | 763.53 | 767.56 | 52.63 | 0.00 |
| α(.), p(sex) | 3 | 763.29 | 769.34 | 54.40 | 0.00 |

POP4

| Model | k | Dev | AICc | Delta | w |
| --- | --- | --- | --- | --- | --- |
| α(t), p(het + sex) | 19 | 9054.58 | 9092.89 | 0.00 | 0.72 |
| α(t + sex), p(het + sex) | 20 | 9054.40 | 9094.74 | 1.85 | 0.28 |
| α(t), p(het) | 18 | 9088.67 | 9124.95 | 32.06 | 0.00 |
| α(t + sex), p(het) | 19 | 9088.66 | 9126.97 | 34.08 | 0.00 |
| α(t), p(sex) | 17 | 9210.51 | 9244.76 | 151.87 | 0.00 |
| α(t + sex), p(sex) | 18 | 9210.49 | 9246.77 | 153.88 | 0.00 |
| α(.), p(het + sex) | 5 | 9254.06 | 9264.08 | 171.19 | 0.00 |
| α(sex), p(het + sex) | 6 | 9253.69 | 9265.73 | 172.84 | 0.00 |
| α(.), p(het) | 4 | 9290.38 | 9298.40 | 205.51 | 0.00 |
| α(t), p(.) | 16 | 9267.24 | 9299.47 | 206.58 | 0.00 |
| α(sex), p(het) | 5 | 9290.38 | 9300.40 | 207.51 | 0.00 |
| α(t + sex), p(.) | 17 | 9267.16 | 9301.41 | 208.52 | 0.00 |
| α(.), p(sex) | 3 | 9400.37 | 9406.38 | 313.49 | 0.00 |
| α(sex), p(sex) | 4 | 9400.27 | 9408.29 | 315.40 | 0.00 |
| α(.), p(.) | 2 | 9458.16 | 9462.16 | 369.27 | 0.00 |
| α(sex), p(.) | 3 | 9458.06 | 9464.07 | 371.18 | 0.00 |

POP5

| Model | k | Dev | AICc | Delta | w |
| --- | --- | --- | --- | --- | --- |
| α(t), p(het + sex) | 23 | 3299.73 | 3346.10 | 0.00 | 0.49 |
| α(t + sex), p(het + sex) | 24 | 3299.50 | 3347.90 | 1.81 | 0.20 |
| α(t), p(sex) | 21 | 3306.98 | 3349.28 | 3.19 | 0.10 |
| α(t + sex), p(het) | 23 | 3303.17 | 3349.54 | 3.44 | 0.09 |
| α(t), p(het) | 22 | 3306.03 | 3350.37 | 4.27 | 0.06 |
| α(t + sex), p(sex) | 22 | 3306.63 | 3350.96 | 4.86 | 0.04 |
| α(t + sex), p(.) | 21 | 3311.35 | 3353.66 | 7.56 | 0.01 |
| α(t), p(.) | 20 | 3314.63 | 3354.91 | 8.81 | 0.01 |
| α(.), p(het + sex) | 5 | 3555.64 | 3565.66 | 219.56 | 0.00 |
| α(.), p(sex) | 3 | 3560.97 | 3566.98 | 220.88 | 0.00 |
| α(sex), p(het + sex) | 6 | 3555.64 | 3567.66 | 221.57 | 0.00 |
| α(sex), p(het) | 5 | 3557.87 | 3567.89 | 221.79 | 0.00 |
| α(sex), p(sex) | 4 | 3560.36 | 3568.37 | 222.27 | 0.00 |
| α(.), p(het) | 4 | 3560.56 | 3568.57 | 222.47 | 0.00 |
| α(sex), p(.) | 3 | 3563.65 | 3569.66 | 223.56 | 0.00 |
| α(.), p(.) | 2 | 3566.53 | 3570.53 | 224.43 | 0.00 |

2. *Examining recruitment at subpopulation level*

In this second part of the appendix, we analyzed the effect of density-dependence and weather variables on recruitment in subpopulations of POP1 (containing four subpopulations: POP1.1, POP1.2, POP1.3, POP1.4) and POP5 (containing four subpopulations: POP5.1, POP5.2, POP5.3, POP5.4). In POP1, the analysis was restricted to POP1.1 and POP1.2 due to the small size of datasets in POP1.3 and POP1.4 (for the number of newt captured in each subpopulation, see Supplementary material S1). In POP5, the analysis was limited to POP5.3 and POP5.4 due to small dataset in POP5.1 and POP5.2. Moreover, in subpopulation POP.5.3, we excluded the first five years of survey (1996-2002) because few newts were captured. We performed the same analyses than those described above.

**Table S4.2.** Modeling recruitment in subpopulations of POP1 (POP1.1 and POP1.2) and POP5 (POP5.3 and POP5.4): selecting the best-supported model without density dependence and weather covariates. α = recruitment, p = recapture, AICc = Akaike information criterion adjusted for small sample size, w = relative AICc weights, k = numbers of parameters.

POP1.1

| Model | k | Dev | AICc | Delta | w |
| --- | --- | --- | --- | --- | --- |
| α(t), p(het) | 21 | 2051.89 | 2094.68 | 0.00 | 0.53 |
| α(t), p(het + sex) | 22 | 2051.75 | 2096.61 | 1.94 | 0.20 |
| α(t + sex), p(het) | 22 | 2051.89 | 2096.75 | 2.07 | 0.19 |
| α(t + sex), p(het + sex) | 23 | 2051.74 | 2098.68 | 4.00 | 0.07 |
| α(t), p(.) | 19 | 2093.02 | 2131.66 | 36.98 | 0.00 |
| α(t), p(sex) | 20 | 2092.97 | 2133.68 | 39.00 | 0.00 |
| α(t + sex), p(.) | 20 | 2092.99 | 2133.70 | 39.03 | 0.00 |
| α(t + sex), p(sex) | 21 | 2092.90 | 2135.68 | 41.01 | 0.00 |
| α(.), p(het) | 4 | 2133.31 | 2141.34 | 46.66 | 0.00 |
| α(.), p(het + sex) | 5 | 2133.21 | 2143.26 | 48.59 | 0.00 |
| α(sex), p(het) | 5 | 2133.30 | 2143.35 | 48.68 | 0.00 |
| α(sex), p(het + sex) | 6 | 2133.20 | 2145.27 | 50.60 | 0.00 |
| α(.), p(.) | 2 | 2186.11 | 2190.12 | 95.44 | 0.00 |
| α(sex), p(.) | 3 | 2186.07 | 2192.09 | 97.41 | 0.00 |
| α(.), p(sex) | 3 | 2186.08 | 2192.10 | 97.42 | 0.00 |
| α(sex), p(sex) | 4 | 2186.00 | 2194.03 | 99.35 | 0.00 |

POP1.2

| Model | k | Dev | AICc | Delta | w |
| --- | --- | --- | --- | --- | --- |
| α(t + sex), p(sex) | 21 | 1133.77 | 1177.40 | 0.00 | 0.90 |
| α(t + sex), p(het + sex) | 23 | 1133.76 | 1181.71 | 4.32 | 0.10 |
| α(t + sex), p(het) | 22 | 1148.72 | 1194.50 | 17.11 | 0.00 |
| α(t), p(het + sex) | 22 | 1157.08 | 1202.86 | 25.46 | 0.00 |
| α(t + sex), p(.) | 20 | 1161.36 | 1202.83 | 25.43 | 0.00 |
| α(t), p(sex) | 20 | 1161.87 | 1203.34 | 25.95 | 0.00 |
| α(t), p(het) | 21 | 1160.85 | 1204.47 | 27.08 | 0.00 |
| α(t), p(.) | 19 | 1170.54 | 1209.87 | 32.47 | 0.00 |
| α(.), p(het) | 4 | 1259.29 | 1267.35 | 89.96 | 0.00 |
| α(sex), p(sex) | 4 | 1259.89 | 1267.96 | 90.56 | 0.00 |
| α(.), p(sex) | 3 | 1262.14 | 1268.19 | 90.79 | 0.00 |
| α(.), p(het + sex) | 5 | 1258.60 | 1268.70 | 91.31 | 0.00 |
| α(sex), p(het) | 5 | 1259.13 | 1269.23 | 91.83 | 0.00 |
| α(sex), p(het + sex) | 6 | 1257.49 | 1269.64 | 92.24 | 0.00 |
| α(sex), p(.) | 3 | 1265.76 | 1271.80 | 94.40 | 0.00 |
| α(.), p(.) | 3 | 1265.84 | 1271.88 | 94.49 | 0.00 |

POP5.3

| Model | k | Dev | AICc | Delta | w |
| --- | --- | --- | --- | --- | --- |
| α(t), p(het + sex) | 17 | 986.07 | 1020.80 | 0.00 | 0.50 |
| α(t + sex), p(het + sex) | 18 | 985.46 | 1022.28 | 1.48 | 0.24 |
| α(t + sex), p(het) | 17 | 988.59 | 1023.32 | 2.52 | 0.14 |
| α(t), p(het) | 16 | 990.90 | 1023.55 | 2.76 | 0.13 |
| α(sex), p(het) | 5 | 1056.15 | 1066.22 | 45.42 | 0.00 |
| α(sex), p(het + sex) | 6 | 1055.43 | 1067.53 | 46.73 | 0.00 |
| α(.), p(het + sex) | 5 | 1058.30 | 1068.37 | 47.57 | 0.00 |
| α(.), p(het) | 4 | 1061.53 | 1069.58 | 48.78 | 0.00 |
| α(t), p(sex) | 15 | 1069.28 | 1099.86 | 79.06 | 0.00 |
| α(t + sex), p(sex) | 16 | 1069.16 | 1101.81 | 81.01 | 0.00 |
| α(t + sex), p(.) | 15 | 1072.10 | 1102.67 | 81.87 | 0.00 |
| α(t), p(.) | 14 | 1074.60 | 1103.11 | 82.31 | 0.00 |
| α(sex), p(.) | 3 | 1141.73 | 1147.76 | 126.96 | 0.00 |
| α(sex), p(sex) | 4 | 1141.73 | 1149.78 | 128.98 | 0.00 |
| α(.), p(sex) | 3 | 1144.46 | 1150.49 | 129.69 | 0.00 |
| α(.), p(.) | 2 | 1146.89 | 1150.90 | 130.10 | 0.00 |

POP5.3

| Model | k | Dev | AICc | Delta | w |
| --- | --- | --- | --- | --- | --- |
| α(t), p(sex) | 21 | 1570.42 | 1613.13 | 0.00 | 0.25 |
| α(t), p(het + sex) | 23 | 1566.49 | 1613.34 | 0.21 | 0.22 |
| α(t), p(het) | 22 | 1568.96 | 1613.74 | 0.61 | 0.18 |
| α(t + sex), p(het + sex) | 24 | 1566.36 | 1615.28 | 2.16 | 0.08 |
| α(t + sex), p(sex) | 22 | 1570.42 | 1615.19 | 2.06 | 0.09 |
| α(t + sex), p(het) | 23 | 1568.45 | 1615.30 | 2.17 | 0.08 |
| α(t), p(.) | 20 | 1575.29 | 1615.94 | 2.81 | 0.06 |
| α(t + sex), p(.) | 21 | 1574.70 | 1617.41 | 4.28 | 0.03 |
| α(.), p(sex) | 3 | 1747.44 | 1753.46 | 140.33 | 0.00 |
| α(.), p(het + sex) | 5 | 1743.46 | 1753.51 | 140.38 | 0.00 |
| α(.), p(het) | 4 | 1746.12 | 1754.15 | 141.02 | 0.00 |
| α(sex), p(het) | 5 | 1745.10 | 1755.15 | 142.02 | 0.00 |
| α(sex), p(sex) | 4 | 1747.32 | 1755.35 | 142.22 | 0.00 |
| α(sex), p(het + sex) | 6 | 1743.46 | 1755.53 | 142.40 | 0.00 |
| α(.), p(.) | 2 | 1752.09 | 1756.10 | 142.97 | 0.00 |
| α (sex), p(.) | 3 | 1750.96 | 1756.98 | 143.85 | 0.00 |

**Table S4.3**. Effect of meteorological variables and density on recruitment in five populations (and two subpopulations) of crested newt. The slope coefficient of each effect and its 95% CI are provided. The results of the ANODEVs (*F*-value and *p*-value) are given and the significant relationships are shown in bold.

|  | TempDFI | TempDFS | RainMM | RainJF | Dens |
| --- | --- | --- | --- | --- | --- |
| **POP1** |  |  |  |  |  |
| Slope | -0.13 | -0.03 | 0.13 | -0.04 | **-0.37** |
| 95% CI | -0.26–-0.003 | -0.16–0.10 | 0.003–0.26 | -0.19–0.10 | **-0.49–0.26** |
| ANODEV | *F* = 0.02, *p* = 0.89 | *F* = 0.46, *p* = 0.51 | *F* = 0.44, *p* = 0.52 | *F* = 0.04, *p* = 0.85 | ***F* = 8.47, *p* = 0.01** |
| **POP1.1** |  |  |  |  |  |
| Slope | -0.22 | -0.14 | 0.14 | 0.11 | **-0.33** |
| 95% CI | -0.38–-0.07 | -0.30–0.02 | -0.02–0.30 | -0.07–0.29 | **-0.48–-0.18** |
| ANODEV | *F* = 1.76, *p* = 0.20 | *F* = 0.61, *p* = 0.45 | *F* = 0.53, *p* = 0.48 | *F* = 0.30, *p* = 0.59 | ***F* = 6.06, *p* = 0.02** |
| **POP1.2** |  |  |  |  |  |
| Slope | 0.11 | 0.35 | 0.07 | -0.49 | -0.96 |
| 95% CI | -0.19–0.41 | -0.01–0.71 | -0.18–0.32 | -0.82–-0.18 | -1.95–0.03 |
| ANODEV | *F* = 0.07, *p* = 0.79 | *F* = 0.61, *p* = 0.44 | *F* = 0.04, *p* = 0.85 | *F* = 1.42, *p* = 0.25 | *F* = 1.44, *p* = 0.25 |
| **POP2** |  |  |  |  |  |
| Slope | 0.05 | -0.004 | 0.10 | 0.17 | 0.15 |
| 95% CI | -0.15–0.26 | -0.21–0.20 | -0.12–0.33 | -0.41–0.07 | -0.37–0.07 |
| ANODEV | *F* = 0.07, *p* = 0.80 | *F* = 0.01, *p* = 0.96 | *F* = 0.20, *p* = 0.67 | *F* = 0.49, *p* = 0.49 | *F* = 0.48, *p* = 0.50 |
| **POP3** |  |  |  |  |  |
| Slope | 0.04 | 0.15 | 0.06 | 0.41 | **0.90** |
| 95% CI | -0.29–0.36 | -0.15–0.45 | -0.17–0.35 | 0.11–0.70 | **0.48–1.33** |
| ANODEV | *F* = 1.73, *p* = 0.24 | *F* = 1.86, *p* = 0.23 | *F* = 1.76, *p* = 0.24 | *F* = 2.93, *p* = 0.15 | ***F* = 8.30, *p* = 0.03** |
| **POP5** |  |  |  |  |  |
| Slope | -0.25 | -0.02 | **-0.38** | -0.26 | 0.26 |
| 95% CI | -0.34–-0.16 | -0.11–0.06 | **-0.47–-0.29** | -0.35–-0.17 | 0.16–0.35 |
| ANODEV | *F* = 2.50, *p* = 0.13 | *F* = 0.02, *p* = 0.87 | ***F* = 7.35, *p* = 0.01** | *F* = 2.69, *p* = 0.12 | *F* = 2.27, *p* = 0.15 |
| **POP5.3** |  |  |  |  |  |
| Slope | -0.53 | **-0.70** | 0.03 | 0.06 | 0.02 |
| 95% CI | -0.81–0.25 | **-0.99–-0.40** | -0.18–0.25 | -0.15–0.27 | -0.19–0.23 |
| ANODEV | *F* = 3.42, *p* = 0.09 | ***F* = 8.18, *p* = 0.01** | *F* = 0.03, *p* = 0.86 | *F* = 0.07, *p* = 0.80 | *F* = 0.02, *p* = 0.88 |
| **POP5.4** |  |  |  |  |  |
| Slope | 0.08 | 0.18 | -0.24 | -0.34 | 0.20 |
| 95% CI | -0.04–0.21 | 0.05–0.31 | -0.36–-0.11 | -0.47–-0.22 | 0.08–0.32 |
| ANODEV | *F* = 0.15, *p* = 0.70 | *F* = 0.79, *p* = 0.39 | *F* = 1.46, *p* = 0.24 | *F* = 3.36, *p* = 0.08 | *F* = 1.21, *p* = 0.29 |
|  | TempMSI | TempMSS | RainAO | RainMS | Dens |
| **POP4** |  |  |  |  |  |
| Slope | **-0.59** | -0.22 | 0.40 | 0.34 | **-0.57** |
| 95% CI | **-0.74–-0.43** | -0.41–-0.05 | 0.27–0.54 | 0.11–0.57 | **-0.71–-0.42** |
| ANODEV | ***F* = 5.54, *p* = 0.03** | *F* = 0.43, *p* = 0.52 | *F* = 2.72, *p* = 0.13 | *F* = 0.60, *p* = 0.45 | ***F* = 5.93, *p* = 0.03** |
