## Appendix S3 for "Drivers of amphibian population dynamics and asynchrony at local and continental scales"

**Appendix 3: Effect of density-dependence and weather on survival**

In this Appendix are provided the model selection procedures performed for survival analyses.

1. *Examining time-specific variation of recapture probabilities in the five populations*

In our analyses, the model [ϕ(t + sex), *p*(het + sex + t)] cannot be run due to recurrent issues of convergence and estimate imprecision. We thus decided to keep [ϕ(t + sex), p(het + sex)] as the most general model because we found recapture heterogeneity (het) in most of the populations (see Table S2.2). Here, we showed that the effect of year (t) was significant (w > 0.9) only in two out of three populations (Table S2.2). For that purpose, we compared the AICc and AICc weight of the model including a yearly-specific recapture probability [ϕ(t), *p*(t)] and the model with a constant recapture probability [ϕ(t), *p*(.)]. In POP1 and POP4, the AICc weight of the model including the year effect was higher than 0.9, indicating significant yearly-specific variation. In these two populations, we then compared the survival estimates provided by [ϕ(t), *p*(t)] and [ϕ(t), *p*(.)] and showed that they do not substantially differ; the mean estimates were relatively similar and the 95% CI greatly overlapped.

**Table S3.1**. Comparing modeling with and without year effect on recapture in five populations of *Triturus cristatus* in Western Europe. ϕ = survival, *p* = recapture, AICc = Akaike information criterion adjusted for small sample size, w = relative AICc weights, k = numbers of parameters, r = rank of the model.

| r | Model | K | Dev. | w | AICc |
| --- | --- | --- | --- | --- | --- |
| **POP1** | |  |  |  |  |
| 1 | ϕ(t), *p*(t) | 35 | 3798.60 | 1.00 | 3869.83 |
| 2 | ϕ(t), *p*(.) | 19 | 3958.85 | 0.00 | 3997.22 |
| **POP2** | |  |  |  |  |
| 1 | ϕ(t), *p*(.) | 31 | 632.37 | 0.58 | 698.62 |
| 2 | ϕ(t), *p*(t) | 17 | 664.04 | 0.41 | 699.31 |
| **POP3** | |  |  |  |  |
| 1 | ϕ(t), *p*(.) | 8 | 484.61 | 0.61 | 500.91 |
| 2 | ϕ(t), *p*(t) | 13 | 475.04 | 0.38 | 501.80 |
| **POP4** | |  |  |  |  |
| 1 | ϕ(t), *p*(t) | 29 | 8968.24 | 1.00 | 9026.95 |
| 2 | ϕ(t), *p*(.) | 15 | 9044.52 | 0.00 | 9074.72 |
| **POP5** | |  |  |  |  |
| 1 | ϕ(t), *p*(.) | 20 | 3243.59 | 1.00 | 3283.87 |
| 2 | ϕ(t), *p*(t) | 37 | 3234.88 | 0.00 | 3309.82 |

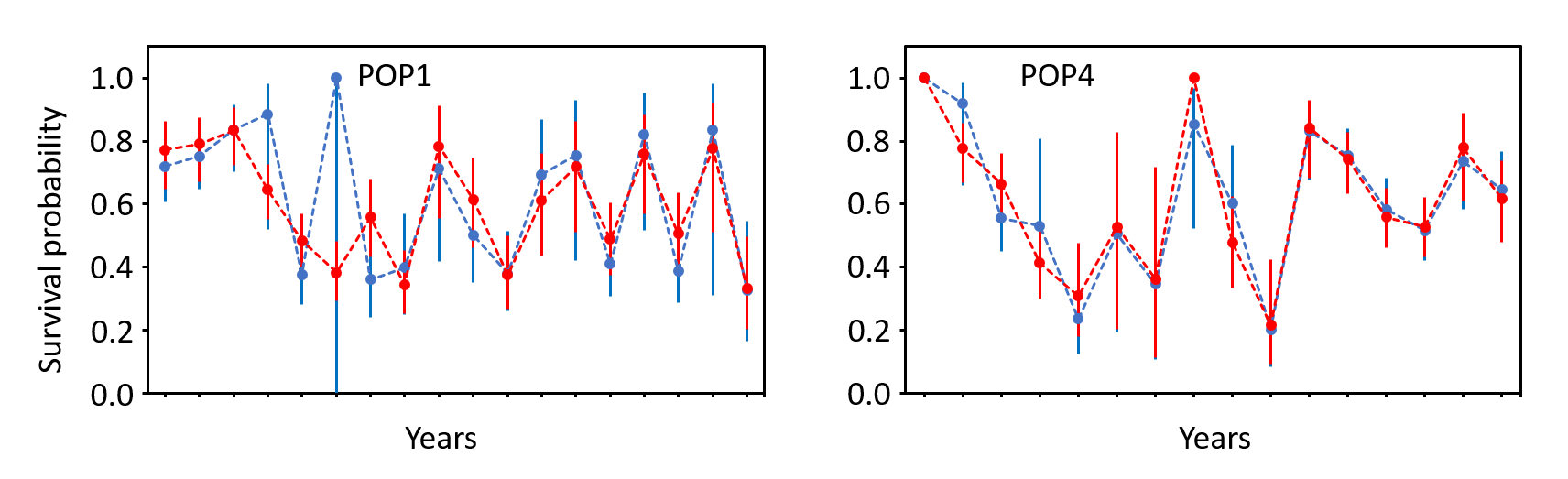

**Fig. S2.1**. Yearly-specific survival probability in POP1 and POP4 estimated using the model [ϕ(t), *p*(t)] (in blue) and the model [ϕ(t), *p*(.)] (in red). 95% CI are shown in error bars.

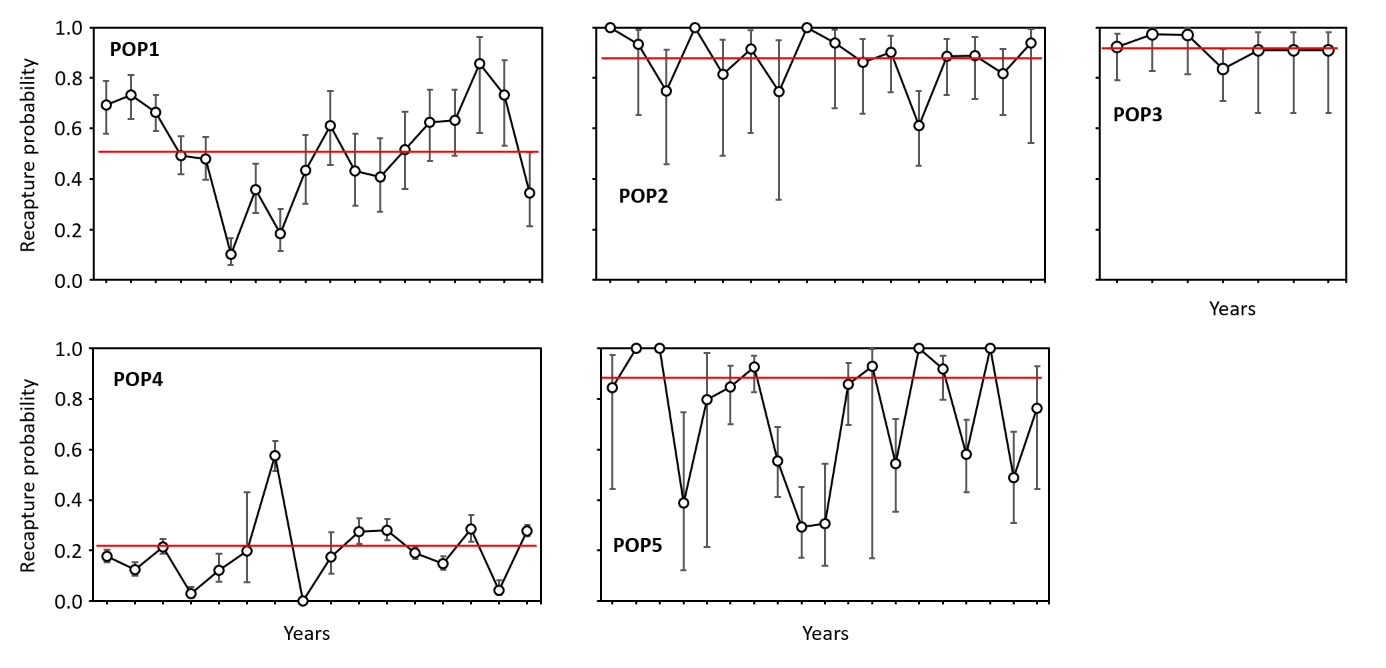

**Fig.2.2.** Time-specific (mean as circles and 95% CI as error bars) and time-constant recapture probability in the five population of *T. cristatus*. The time-specific recapture probability is extracted from the model φ(.), p(t) and time-constant recapture probability from φ(.), p(.).

2. *Examining survival at the population level*

In this second part of the appendix, we provide analyses carried out separately for each of the five studied populations of *Triturus cristatus* (POP1, POP2, POP3, POP4 and POP5). First, for each population, we defined the best-supported model without weather covariate using Akaile information criteria adjusted for small sample size (AICc) and relative AICc weights (**Table S2.1**). We first modeled recapture and tested the hypotheses that recapture varied according heterogeneity groups (‘het’) and between genders (‘sex’). Then, retaining the best combination of effects on recapture, we modeled survival and examined the hypotheses that survival varied between years (‘t’) and genders (‘sex’).

**Table S3.2**. POP1. Survival and recapture in five populations of *Triturus cristatus* in Western Europe: model selection procedure. ϕ = survival, p = recapture, AICc = Akaike information criterion adjusted for small sample size, w = relative AICc weights, k = numbers of parameters.

| Model | k | Dev | AICc | Delta | w |
| --- | --- | --- | --- | --- | --- |
| ϕ(t + sex), p(het + sex) | 23 | 3860.57 | 3907.11 | 0.00 | 0.84 |
| ϕ(t), p(het + sex) | 22 | 3865.91 | 3910.40 | 3.29 | 0.16 |
| ϕ(t), p(het) | 21 | 3876.44 | 3918.89 | 11.78 | 0.00 |
| ϕ(t + sex), p(het) | 22 | 3876.27 | 3920.76 | 13.65 | 0.00 |
| ϕ(t + sex), p(sex) | 21 | 3941.98 | 3984.43 | 77.32 | 0.00 |
| ϕ(t), p(sex) | 20 | 3944.53 | 3984.94 | 77.83 | 0.00 |
| ϕ(t), p(.) | 19 | 3958.85 | 3997.22 | 90.11 | 0.00 |
| ϕ(t + sex), p(.) | 20 | 3958.81 | 3999.21 | 92.11 | 0.00 |
| ϕ(.), p(het + sex) | 5 | 4049.39 | 4059.42 | 152.31 | 0.00 |
| ϕ(sex), p(het + sex) | 6 | 4048.48 | 4060.52 | 153.41 | 0.00 |
| ϕ(.), p(het) | 4 | 4056.95 | 4064.97 | 157.86 | 0.00 |
| ϕ(sex), p(het) | 5 | 4056.58 | 4066.61 | 159.50 | 0.00 |
| ϕ(.), p(sex) | 3 | 4111.83 | 4117.84 | 210.73 | 0.00 |
| ϕ(sex), p(sex) | 4 | 4111.66 | 4119.68 | 212.57 | 0.00 |
| ϕ(.), p(.) | 2 | 4122.95 | 4126.95 | 219.85 | 0.00 |
| ϕ(sex), p(.) | 3 | 4122.17 | 4128.18 | 221.07 | 0.00 |

POP2

| Model | k | Dev | AICc | Delta | w |
| --- | --- | --- | --- | --- | --- |
| ϕ(.), p(het + sex) | 5 | 675.07 | 685.19 | 0.00 | 0.31 |
| ϕ(.), p(sex) | 3 | 679.48 | 685.53 | 0.34 | 0.26 |
| ϕ(sex), p(sex) | 4 | 678.59 | 686.67 | 1.48 | 0.15 |
| ϕ(sex), p(het + sex) | 6 | 674.61 | 686.78 | 1.59 | 0.14 |
| ϕ(t), p(sex) | 18 | 652.15 | 689.57 | 4.39 | 0.03 |
| ϕ(t), p(het + sex) | 20 | 648.34 | 690.10 | 4.91 | 0.03 |
| ϕ(t + sex), p(sex) | 19 | 650.98 | 690.56 | 5.37 | 0.02 |
| ϕ(t + sex), p(het + sex) | 21 | 647.67 | 691.61 | 6.42 | 0.01 |
| ϕ(.), p(het) | 4 | 682.11 | 690.19 | 5.00 | 0.03 |
| ϕ(sex), p(het) | 5 | 680.96 | 691.08 | 5.89 | 0.02 |
| ϕ(t), p(het) | 19 | 655.84 | 695.43 | 10.24 | 0.00 |
| ϕ(t + sex), p(het) | 20 | 654.27 | 696.02 | 10.83 | 0.00 |
| ϕ(.), p(.) | 2 | 691.28 | 695.31 | 10.12 | 0.00 |
| ϕ(sex), p(.) | 3 | 689.47 | 695.52 | 10.33 | 0.00 |
| ϕ(t + sex), p(.) | 18 | 661.68 | 699.10 | 13.91 | 0.00 |
| ϕ(t), p(.) | 17 | 664.04 | 699.31 | 14.13 | 0.00 |

POP3

| Model | k | Dev | AICc | Delta | w |
| --- | --- | --- | --- | --- | --- |
| ϕ(.), p(het + sex) | 5 | 466.14 | 476.26 | 0.00 | 0.30 |
| ϕ(.), p(het) | 4 | 468.22 | 476.30 | 0.04 | 0.30 |
| ϕ(sex), p(het + sex) | 6 | 464.52 | 476.70 | 0.44 | 0.24 |
| ϕ(sex), p(het) | 5 | 467.67 | 477.80 | 1.54 | 0.14 |
| ϕ(t), p(het) | 10 | 461.62 | 482.07 | 5.81 | 0.02 |
| ϕ(t + sex), p(het) | 11 | 461.44 | 483.99 | 7.73 | 0.01 |
| ϕ(.), p(.) | 2 | 487.60 | 491.63 | 15.37 | 0.00 |
| ϕ(sex), p(.) | 3 | 486.38 | 492.43 | 16.17 | 0.00 |
| ϕ(.), p(sex) | 3 | 487.59 | 493.64 | 17.38 | 0.00 |
| ϕ(t), p(het + sex) | 11 | 471.69 | 494.24 | 17.98 | 0.00 |
| ϕ(sex), p(sex) | 4 | 486.28 | 494.36 | 18.10 | 0.00 |
| ϕ(t + sex), p(het + sex) | 12 | 470.90 | 495.55 | 19.29 | 0.00 |
| ϕ(t), p(.) | 8 | 484.61 | 500.91 | 24.65 | 0.00 |
| ϕ(t + sex), p(.) | 9 | 483.47 | 501.84 | 25.58 | 0.00 |
| ϕ(t), p(sex) | 9 | 484.59 | 502.97 | 26.71 | 0.00 |
| ϕ(t + sex), p(sex) | 10 | 483.36 | 503.82 | 27.56 | 0.00 |

POP4

| Model | k | Dev | AICc | Delta | w |
| --- | --- | --- | --- | --- | --- |
| ϕ(t), p(het + sex) | 18 | 8929.67 | 8965.95 | 0.00 | 0.73 |
| ϕ(t + sex), p(het + sex) | 19 | 8929.64 | 8967.95 | 2.01 | 0.27 |
| ϕ(t), p(het) | 17 | 8944.14 | 8978.39 | 12.44 | 0.00 |
| ϕ(t + sex), p(het) | 18 | 8944.10 | 8980.38 | 14.43 | 0.00 |
| ϕ(t), p(sex) | 16 | 9019.88 | 9052.10 | 86.15 | 0.00 |
| ϕ(t + sex), p(sex) | 17 | 9019.87 | 9054.12 | 88.17 | 0.00 |
| ϕ(t), p(.) | 15 | 9044.52 | 9074.72 | 108.77 | 0.00 |
| ϕ(t + sex), p(.) | 16 | 9044.31 | 9076.53 | 110.58 | 0.00 |
| ϕ(.), p(het + sex) | 5 | 9116.00 | 9126.03 | 160.08 | 0.00 |
| ϕ(sex), p(het + sex) | 6 | 9115.75 | 9127.79 | 161.84 | 0.00 |
| ϕ(.), p(het) | 4 | 9129.65 | 9137.67 | 171.72 | 0.00 |
| ϕ(sex), p(het) | 5 | 9129.62 | 9139.65 | 173.70 | 0.00 |
| ϕ(.), p(sex) | 3 | 9206.68 | 9212.69 | 246.75 | 0.00 |
| ϕ(sex), p(sex) | 4 | 9206.51 | 9214.52 | 248.58 | 0.00 |
| ϕ(.), p(.) | 2 | 9229.76 | 9233.77 | 267.82 | 0.00 |
| ϕ(sex), p(.) | 3 | 9229.76 | 9235.77 | 269.82 | 0.00 |

POP5

| Model | k | Dev | AICc | Delta | w |
| --- | --- | --- | --- | --- | --- |
| ϕ(t), p(het) | 22 | 3233.12 | 3277.46 | 0.00 | 0.39 |
| ϕ(t + sex), p(het) | 23 | 3232.26 | 3278.63 | 1.17 | 0.22 |
| ϕ(t), p(het + sex) | 23 | 3232.33 | 3278.70 | 1.25 | 0.21 |
| ϕ(t + sex), p(het + sex) | 24 | 3231.32 | 3279.72 | 2.26 | 0.13 |
| ϕ(t), p(sex) | 21 | 3240.80 | 3283.11 | 5.65 | 0.02 |
| ϕ(t), p(.) | 20 | 3243.59 | 3283.87 | 6.41 | 0.02 |
| ϕ(t + sex), p(sex) | 22 | 3240.79 | 3285.13 | 7.67 | 0.01 |
| ϕ(t + sex), p(.) | 21 | 3242.86 | 3285.17 | 7.71 | 0.01 |
| ϕ(.), p(het + sex) | 5 | 3432.68 | 3442.70 | 165.24 | 0.00 |
| ϕ(.), p(sex) | 3 | 3436.71 | 3442.72 | 165.27 | 0.00 |
| ϕ(sex), p(sex) | 4 | 3436.37 | 3444.39 | 166.93 | 0.00 |
| ϕ(sex), p(het + sex) | 6 | 3432.67 | 3444.70 | 167.24 | 0.00 |
| ϕ(.), p(het) | 4 | 3437.05 | 3445.07 | 167.61 | 0.00 |
| ϕ(sex), p(het) | 5 | 3435.17 | 3445.19 | 167.73 | 0.00 |
| ϕ(.), p(.) | 2 | 3441.53 | 3445.53 | 168.08 | 0.00 |
| ϕ(sex), p(.) | 3 | 3439.55 | 3445.55 | 168.10 | 0.00 |

3. *Examining survival at subpopulation level*

In this second part of the appendix, we analyzed the effect of density-dependence and weather variables on survival in subpopulations of POP1 (containing four subpopulations: POP1.1, POP1.2, POP1.3, POP1.4) and POP5 (containing four subpopulations: POP5.1, POP5.2, POP5.3, POP5.4). In POP1, the analysis was restricted to POP1.1 and POP1.2 due to the reduced size of datasets in POP1.3 and POP1.4 (for the number of newt captured in each subpopulation, see Supplementary material S1). In POP5, the analysis was limited to POP5.3 and POP5.4 due to small dataset in POP5.1 and POP5.2. Moreover, in subpopulation POP.5.3, we excluded the first five years of survey (1996-2002) because few newts were captured. We performed the same analyses than those described above.

**Table S3.3.** Modeling survival in subpopulations of POP1 (POP1.1 and POP1.2) and POP5 (POP5.3 and POP5.4): selecting the best-supported model without density dependence and weather covariates. ϕ = survival, *p* = recapture, AICc = Akaike information criterion adjusted for small sample size, w = relative AICc weights, k = numbers of parameters.

POP1.1

| Model | k | Dev | AICc | Delta | w |
| --- | --- | --- | --- | --- | --- |
| ϕ(t), p(het) | 21 | 2013.74 | 2056.52 | 0.00 | 0.54 |
| ϕ(t + sex), p(het) | 22 | 2013.71 | 2058.56 | 2.04 | 0.20 |
| ϕ(t), p(het + sex) | 22 | 2013.71 | 2058.57 | 2.05 | 0.19 |
| ϕ(t + sex), p(het + sex) | 23 | 2013.69 | 2060.63 | 4.11 | 0.07 |
| ϕ(t), p(.) | 19 | 2075.60 | 2114.24 | 57.71 | 0.00 |
| ϕ(t + sex), p(.) | 20 | 2075.58 | 2116.29 | 59.76 | 0.00 |
| ϕ(t), p(sex) | 20 | 2075.59 | 2116.30 | 59.78 | 0.00 |
| ϕ(t + sex), p(sex) | 21 | 2084.28 | 2127.06 | 70.54 | 0.00 |
| ϕ(.), p(het) | 4 | 2161.97 | 2170.00 | 113.48 | 0.00 |
| ϕ(.), p(het + sex) | 5 | 2161.88 | 2171.93 | 115.41 | 0.00 |
| ϕ(sex), p(het) | 5 | 2161.94 | 2171.99 | 115.46 | 0.00 |
| ϕ(sex), p(het + sex) | 6 | 2161.80 | 2173.87 | 117.34 | 0.00 |
| ϕ(.), p(.) | 2 | 2202.89 | 2206.90 | 150.38 | 0.00 |
| ϕ(sex), p(.) | 3 | 2202.78 | 2208.80 | 152.27 | 0.00 |
| ϕ(.), p(sex) | 3 | 2202.88 | 2208.90 | 152.38 | 0.00 |
| ϕ(sex), p(sex) | 4 | 2202.73 | 2210.76 | 154.24 | 0.00 |

POP1.2

| Model | k | Dev | AICc | Delta | w |
| --- | --- | --- | --- | --- | --- |
| ϕ(t + sex), p(sex) | 21 | 1144,26 | 1187,88 | 0,00 | 0,65 |
| ϕ(t + sex), p(het + sex) | 23 | 1141,17 | 1189,12 | 1,23 | 0,35 |
| ϕ(t), p(sex) | 20 | 1161,08 | 1202,55 | 14,67 | 0,00 |
| ϕ(t), p(het + sex) | 22 | 1158,38 | 1204,16 | 16,28 | 0,00 |
| ϕ(t + sex), p(het) | 22 | 1160,15 | 1205,93 | 18,05 | 0,00 |
| ϕ(t + sex), p(.) | 20 | 1165,93 | 1207,40 | 19,52 | 0,00 |
| ϕ(t), p(het) | 21 | 1165,49 | 1209,12 | 21,24 | 0,00 |
| ϕ(t), p(.) | 19 | 1170,44 | 1209,77 | 21,89 | 0,00 |
| ϕ(.), p(het) | 4 | 1272,70 | 1280,77 | 92,89 | 0,00 |
| ϕ(.), p(sex) | 3 | 1275,31 | 1281,35 | 93,47 | 0,00 |
| ϕ(.), p(het + sex) | 5 | 1271,64 | 1281,75 | 93,87 | 0,00 |
| ϕ(sex), p(sex) | 4 | 1274,28 | 1282,35 | 94,47 | 0,00 |
| ϕ(sex), p(het) | 5 | 1272,70 | 1282,80 | 94,92 | 0,00 |
| ϕ(sex), p(het + sex) | 6 | 1271,27 | 1283,41 | 95,53 | 0,00 |
| ϕ(.), p(.) | 2 | 1279,70 | 1283,72 | 95,84 | 0,00 |
| ϕ(sex), p(.) | 3 | 1279,68 | 1285,72 | 97,84 | 0,00 |

POP5.3

| Model | k | Dev | AICc | Delta | w |
| --- | --- | --- | --- | --- | --- |
| ϕ(t), p(.) | 14 | 788,53 | 817,03 | 0,00 | 0,24 |
| ϕ(t), p(het) | 16 | 784,88 | 817,53 | 0,51 | 0,19 |
| ϕ(t + sex), p(.) | 15 | 787,39 | 817,96 | 0,94 | 0,15 |
| ϕ(t + sex), p(het) | 17 | 783,65 | 818,39 | 1,36 | 0,12 |
| ϕ(t), p(het + sex) | 17 | 783,75 | 818,49 | 1,46 | 0,12 |
| ϕ(t), p(sex) | 15 | 788,46 | 819,03 | 2,00 | 0,09 |
| ϕ(t + sex), p(sex) | 16 | 787,31 | 819,96 | 2,93 | 0,06 |
| ϕ(t + sex), p(het + sex) | 18 | 783,49 | 820,31 | 3,28 | 0,05 |
| ϕ(sex), p(.) | 3 | 875,58 | 881,61 | 64,59 | 0,00 |
| ϕ(.), p(.) | 2 | 878,60 | 882,62 | 65,59 | 0,00 |
| ϕ(sex), p(sex) | 4 | 875,57 | 883,62 | 66,60 | 0,00 |
| ϕ(.), p(het + sex) | 5 | 874,20 | 884,27 | 67,24 | 0,00 |
| ϕ(.), p(sex) | 3 | 878,23 | 884,26 | 67,23 | 0,00 |
| ϕ(sex), p(het) | 5 | 875,26 | 885,33 | 68,30 | 0,00 |
| ϕ(sex), p(het + sex) | 6 | 874,16 | 886,26 | 69,24 | 0,00 |
| ϕ(.), p(het) | 4 | 878,21 | 886,26 | 69,23 | 0,00 |

POP5.4

| Model | k | Dev | AICc | Delta | w |
| --- | --- | --- | --- | --- | --- |
| ϕ(t), p(het) | 22 | 1591,3526 | 1636,1287 | 0,00 | 0,28 |
| ϕ(t), p(sex) | 21 | 1594,1128 | 1636,8208 | 0,69 | 0,20 |
| ϕ(t), p(het + sex) | 23 | 1590,2156 | 1637,0629 | 0,93 | 0,17 |
| ϕ(t + sex), p(het) | 23 | 1590,9616 | 1637,8089 | 1,68 | 0,12 |
| ϕ(t + sex), p(sex) | 22 | 1594,0992 | 1638,8753 | 2,75 | 0,07 |
| ϕ(t), p(.) | 20 | 1598,201 | 1638,8442 | 2,72 | 0,07 |
| ϕ(t + sex), p(het + sex) | 24 | 1590,2087 | 1639,1303 | 3,00 | 0,06 |
| ϕ(t + sex), p(.) | 21 | 1597,7722 | 1640,4802 | 4,35 | 0,03 |
| ϕ(.), p(sex) | 3 | 1710,2882 | 1716,3063 | 80,18 | 0,00 |
| ϕ(.), p(het + sex) | 5 | 1707,0065 | 1717,0519 | 80,92 | 0,00 |
| ϕ(.), p(het) | 4 | 1709,2982 | 1717,3285 | 81,20 | 0,00 |
| ϕ(sex), p(sex) | 4 | 1710,2858 | 1718,3161 | 82,19 | 0,00 |
| ϕ(.), p(.) | 2 | 1714,3615 | 1718,3705 | 82,24 | 0,00 |
| ϕ(sex), p(het) | 5 | 1708,8651 | 1718,9105 | 82,78 | 0,00 |
| ϕ(sex), p(het + sex) | 6 | 1706,9782 | 1719,0418 | 82,91 | 0,00 |
| ϕ(sex), p(.) | 3 | 1713,8688 | 1719,887 | 83,76 | 0,00 |

**Table S3.4**. Effect of meteorological variables and density on survival in five populations (and two subpopulations) of crested newt. The slope coefficient of each effect and its 95% CI are provided. The results of the ANODEVs (*F*-value and *p*-value) are given and the significant relationships are shown in bold.

|  | TempDFI | TempDFS | RainMM | RainJF | Dens |
| --- | --- | --- | --- | --- | --- |
| **POP1** |  |  |  |  |  |
| Slope | -0.001 | -0.25 | **-0.62** | **-0.50** | 0.16 |
| 95% CI | -0.12–0.12 | -0.38–-0.12 | **-0.64–-0.41** | **-0.65–-0.35** | 0.07–0.25 |
| ANODEV | *F* = 0.001, *p* = 0.98 | *F* = 1.49, *p* = 0.24 | ***F* = 12.97, *p* = 0.002** | ***F* = 5.64, *p* = 0.03** | *F* = 1.11, *p* = 0.31 |
| **POP1.1** |  |  |  |  |  |
| Slope | -0.10 | -0.28 | **-0.62** | **-0.54** | -0.25 |
| 95% CI | -0.25–0.05 | -0.44–-0.11 | **-0.76–-0.45** | **-0.74–-0.35** | -0.36–-0.13 |
| ANODEV | *F* = 0.19, *p* = 0.67 | *F* = 1.39, *p* = 0.26 | ***F* = 11.68, *p* = 0.003** | ***F* = 5.14, *p* = 0.04** | *F* = 2.56, *p* = 0.14 |
| **POP1.2** |  |  |  |  |  |
| Slope | 0.45 | -0.07 | -0.37 | -0.25 | **-0.57** |
| 95% CI | 0.20–0.69 | -0.32–0.18 | -0.57–-0.16 | -0.55–0.04 | **-0.81–-0.35** |
| ANODEV | *F* = 1.69, *p* = 0.21 | *F* = 0.04, *p* = 0.84 | *F* = 1.45, *p* = 0.24 | *F* = 0.38, *p* = 0.55 | ***F* = 4.73, *p* = 0.04** |
| **POP2** |  |  |  |  |  |
| Slope | -0.13 | -0.14 | -0.14 | -0.33 | 0.01 |
| 95% CI | -0.39–0.13 | -0.40–0.12 | -0.41–0.12 | -0.67–0.01 | -0.28–0.29 |
| ANODEV | *F* = 0.50, *p* = 0.49 | *F* = 0.60, *p* = 0.45 | *F* = 0.58, *p* = 0.46 | *F* = 2.39, *p* = 0.14 | *F* = 0.005, *p* = 0.94 |
| **POP3** |  |  |  |  |  |
| Slope | -0.24 | -0.25 | 0.03 | -0.10 | 0.14 |
| 95% CI | -0.79–0.32 | -0.84–0.34 | -0.36–0.42 | -0.58–0.39 | -0.78–0.49 |
| ANODEV | *F* = 0.96, *p* = 0.37 | *F* = 0.94, *p* = 0.38 | *F* = 0.15, *p* = 0.72 | *F* = 0.20, *p* = 0.68 | *F* = 0.24, *p* = 0.64 |
| **POP5** |  |  |  |  |  |
| Slope | -0.15 | -0.16 | 0.03 | 0.02 | 0.08 |
| 95% CI | -0.24–-0.07 | -0.26–-0.06 | -0.05–0.11 | -0.09–0.12 | -0.03–0.18 |
| ANODEV | *F* = 1.14, *p* = 0.30 | *F* = 0.96, *p* = 0.34 | *F* = 0.04, *p* = 0.85 | *F* = 0.01, *p* = 0.93 | *F* = 0.18, *p* = 0.67 |
| **POP5.3** |  |  |  |  |  |
| Slope | 0.17 | 0.15 | 0.03 | -0.05 | 0.08 |
| 95% CI | -0.01–0.36 | -0.04–0.33 | -0.39–0.15 | -0.28–0.16 | -0.04–0.17 |
| ANODEV | *F* = 0.44, *p* = 0.52 | *F* = 0.31, *p* = 0.59 | *F* = 0.55, *p* = 0.47 | *F* = 0.01, *p* = 0.86 | *F* = 0.06, *p* = 0.82 |
| **POP5.4** |  |  |  |  |  |
| Slope | 0.11 | 0.13 | -0.13 | -0.05 | 0.21 |
| 95% CI | -0.03–0.26 | -0.01–0.27 | -0.26–-0.001 | -0.19–0.08 | 0.08–0.33 |
| ANODEV | *F* = 0.38, *p* = 0.54 | *F* = 0.55, *p* = 0.47 | *F* = 0.66, *p* = 0.42 | *F* = 0.10, *p* = 0.75 | *F* = 1.96, *p* = 0.17 |
|  | TempMSI | TempMSS | RainAO | RainMS | Dens |
| **POP4** |  |  |  |  |  |
| Slope | 0.15 | 0.02 | 0.49 | **-0.57** | 0.14 |
| 95% CI | -0.08–0.37 | -0.22–0.26 | 0.29–0.26 | **-0.71–-0.42** | -0.01–0.31 |
| ANODEV | *F* = 0.03, *p* = 0.96 | *F* = 0.11, *p* = 0.74 | *F* = 1.93, *p* = 0.19 | ***F* = 5.91, *p* = 0.03** | *F* = 0.22, *p* = 0.65 |
