## Appendix S5 for "Drivers of amphibian population dynamics and asynchrony at local and continental scales"

**Appendix 5: Effect of density-dependence and local weather on population growth rate**

In this Appendix, we provide the outputs of Gompertz state-space (GSS) models used to examine the influence of density dependence and weather variables on population growth rate. We first analyzed the effect of population density and local weather variation in the five studied populations. Then, we examined these effects within population POP1 and POP5.

1. *Examining demographic growth rate at the population level*

First, we analyzed the effect of population density at *t* – 1 on population growth rate. The table **S5.1** and figure **S5.1** provided below shows the outputs of thee GSS models in which we tested the effect of population density at *t* – 1.

**Table S5.1.** Effect of population density at *t* – 1 on the population growth rate in five populations of *Triturus cristatus* in Western Europe. The table shows the estimates (with their standard deviation in brackets) and 95% credible intervals of the parameters for the GSS model. $\sigma_{obs}^{^{2}}$ = the observation variance, $\sigma_{proc}^{^{2}}$ = the process variance, *a* = intrinsic rate of increase, $b_{1}$ = the coefficient slope for density at *t* – 1.

| Populations | Parameter | Estimate (sd) | 2.5% | 97.5% |
| --- | --- | --- | --- | --- |
| **POP1** | $\sigma_{proc}^{^{2}}$ | 0.13 (0.08) | 0.01 | 0.33 |
|  | $\sigma_{obs}^{^{2}}$ | 0.05 (0.06) | 0.00 | 0.22 |
|  | *a* | 1.44 (1.28) | -0.58 | 4.11 |
|  | $b_{1}$ | -0.28 (0.24) | -0.79 | 0.10 |
| **POP2** | $\sigma_{proc}^{^{2}}$ | 0.12 (0.09) | 0.01 | 0.35 |
|  | $\sigma_{obs}^{^{2}}$ | 0.05 (0.05) | 0.00 | 0.18 |
|  | *a* | 0.30 (0.60) | -0.76 | 1.62 |
|  | $b_{1}$ | -0.06 (0.18) | -0.45 | 0.25 |
| **POP3** | $\sigma_{proc}^{^{2}}$ | 0.25 (0.86) | 0.00 | 1.39 |
|  | $\sigma_{obs}^{^{2}}$ | 0.19 (0.74) | 0.00 | 1.12 |
|  | *a* | 1.85 (1.17) | 0.00 | 4.58 |
|  | $b_{1}$ | -0.39 (0.30) | -1.12 | 0.11 |
| **POP4** | $\sigma_{proc}^{^{2}}$ | 2.30 (2.31) | 0.01 | 8.05 |
|  | $\sigma_{obs}^{^{2}}$ | 2.53 (2.29) | 0.01 | 8.13 |
|  | *a* | 3.58 (1.97) | 0.20 | 7.88 |
|  | $b_{1}$ | -0.90 (0.46) | -1.86 | -0.07 |
| **POP5** | $\sigma_{proc}^{^{2}}$ | 0.20 (0.17) | 0.00 | 0.60 |
|  | $\sigma_{obs}^{^{2}}$ | 0.15 (0.15) | 0.00 | 0.51 |
|  | *a* | 2.11 (1.16) | 0.26 | 4.72 |
|  | $b_{1}$ | -0.41 (0.23) | -0.94 | -0.04 |

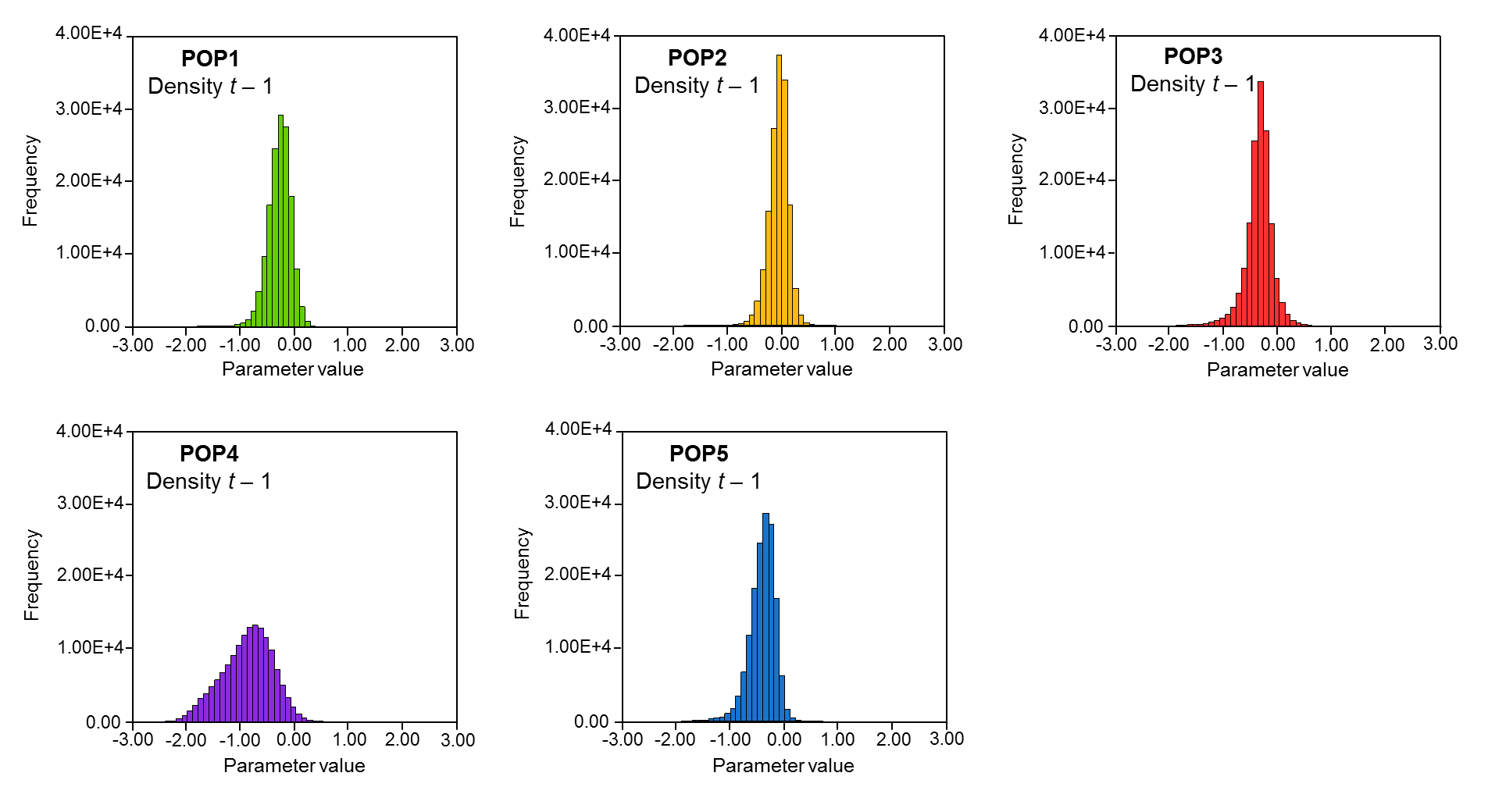

**Fig. S5.1.** Effect of population density at *t* – 1 on the population growth rate in five populations of *Triturus cristatus* in Western Europe. The figure shows the distribution of posterior estimates for density effects at *t* – 1.

Then, we examined the influence of local weather variation on population growth rate. The tables **S5.2** and **S5.3** and the figure S4.2 provided below shows the outputs of thee GSS models in which we tested the effect of population density at *t* – 1. In POP1, POP2, POP3 and POP5, the four following weather variables were considered in our analyses: the cumulative rainfall during the activity period (March-May) (‘rainMM’) and the non-activity period (June-February) (‘rainJF’); the minimum (‘tempDFI’) and maximum (‘tempDFS’) monthly temperature during winter (December-February). In POP4, the four following weather variables were considered: the cumulative rainfall during the activity period (October-April) (‘rainAO’) and the non-activity period (May-September) (‘rainMS’); the minimum (‘tempMSI’) and maximum (‘tempMSS’) monthly temperature during the non-activity period (May-September).

**Table S5.2.** Effects of temperature on the population growth rate in five populations of *Triturus cristatus* in Western Europe. The table shows the estimates (with their standard deviation in brackets) and 95% credible intervals of the parameters for the GSS models including the local minimum and maximum temperature. $\sigma_{obs}^{^{2}}$ = the observation variance, $\sigma_{proc}^{^{2}}$ = the process variance, *a* = intrinsic rate of increase, $b_{2}$ = le slope for the temperature effect.

|  |  | Min. temperature  (tempDFI, tempMSI) | | | Max. temperature  (tempDFS, tempMSS) | | |
| --- | --- | --- | --- | --- | --- | --- | --- |
| Populations | Parameter | Estimate (sd) | 2.5% | 97.5% | Estimate (sd) | 2.5% | 97.5% |
| **POP1** | $\sigma_{proc}^{^{2}}$ | 0.11 (0.09) | 0.00 | 0.34 | 0.13 (0.10) | 0.00 | 0.36 |
|  | $\sigma_{obs}^{^{2}}$ | 0.06 (0.06) | 0.00 | 0.20 | 0.05 (0.06) | 0.00 | 0.21 |
|  | *a* | -0.04 (0.08) | -0.20 | 0.13 | -0.04 (0.09) | -0.21 | 0.15 |
|  | $b_{2}$ | -0.05 (0.10) | -0.24 | 0.15 | 0.00 (0.12) | -0.22 | 0.23 |
| **POP2** | $\sigma_{proc}^{^{2}}$ | 0.11 (0.09) | 0.00 | 0.34 | 0.11 (0.09) | 0.00 | 0.33 |
|  | $\sigma_{obs}^{^{2}}$ | 0.06 (0.05) | 0.00 | 0.19 | 0.05 (0.05) | 0.00 | 0.19 |
|  | *a* | 0.09 (0.08) | -0.08 | 0.26 | 0.09 (0.08) | -0.09 | 0.26 |
|  | $b_{2}$ | 0.02 (0.10) | -0.18 | 0.21 | -0.05 (0.11) | -0.27 | 0.16 |
| **POP3** | $\sigma_{proc}^{^{2}}$ | 0.50 (1.78) | 0.00 | 2.82 | 0.45 (1.57) | 0.00 | 2.52 |
|  | $\sigma_{obs}^{^{2}}$ | 0.24 (1.00) | 0.00 | 1.35 | 0.22 (0.86) | 0.00 | 1.17 |
|  | *a* | 0.40 (0.29) | -0.14 | 0.97 | 0.41 (0.27) | -0.10 | 0.93 |
|  | $b_{2}$ | -0.20 (0.40) | -0.95 | 0.58 | -0.22 (0.35) | -0.88 | 0.46 |
| **POP4** | $\sigma_{proc}^{^{2}}$ | 2.52 (2.48) | 0.01 | 8.71 | 3.47 (3.02) | 0.05 | 9.95 |
|  | $\sigma_{obs}^{^{2}}$ | 1.61 (1.53) | 0.01 | 5.39 | 1.81 (1.90) | 0.01 | 6.64 |
|  | *a* | -0.10 (0.43) | -1.01 | 0.79 | 0.07 (0.49) | -0.96 | 1.08 |
|  | $b_{2}$ | -1.11 (0.63) | -2.39 | 0.12 | -0.56 (0.70) | -1.94 | 0.83 |
| **POP5** | $\sigma_{proc}^{^{2}}$ | 0.17 (0.19) | 0.00 | 0.65 | 0.22 (0.21) | 0.00 | 0.74 |
|  | $\sigma_{obs}^{^{2}}$ | 0.17 (0.12) | 0.00 | 0.46 | 0.16 (0.13) | 0.00 | 0.48 |
|  | *a* | 0.10 (1.10) | -0.11 | 0.30 | 0.10 (0.11) | -0.14 | 0.33 |
|  | $b_{2}$ | 0.17 (0.13) | -0.12 | 0.44 | 0.08 (0.14) | -0.22 | 0.13 |

**Table S5.3.** Effect of rainfall on the population growth rate in five populations of *Triturus cristatus* in Western Europe. The table shows the estimates (with their standard deviation in brackets) and 95% credible intervals of the parameters for the GSS models including the local cumulative rainfall. $\sigma_{obs}^{^{2}}$ = the observation variance, $\sigma_{proc}^{^{2}}$ = the process variance, *a* = intrinsic rate of increase, $b_{2}$ = le slope for the rainfall effect.

|  |  | Rainfall (rainMM, rainOA) | | | Rainfall (rainJF, rainMS) | | |
| --- | --- | --- | --- | --- | --- | --- | --- |
| Populations | Parameter | Estimate (sd) | 2.5% | 97.5% | Estimate (sd) | 2.5% | 97.5% |
| **POP1** | $\sigma_{proc}^{^{2}}$ | 0.11 (0.08) | 0.00 | 0.31 | 0.11 (0.08) | 0.00 | 0.32 |
|  | $\sigma_{obs}^{^{2}}$ | 0.05 (0.05) | 0.00 | 0.18 | 0.05 (0.05) | 0.00 | 0.18 |
|  | *a* | -0.04 (0.08) | -0.20 | 0.13 | -0.04 (0.08) | -0.20 | 0.13 |
|  | $b_{2}$ | -0.13 (0.10) | -0.33 | 0.06 | -0.12 (0.09) | -0.31 | 0.07 |
| **POP2** | $\sigma_{proc}^{^{2}}$ | 0.09 (0.09) | 0.00 | 0.31 | 0.10 (0.09) | 0.00 | 0.31 |
|  | $\sigma_{obs}^{^{2}}$ | 0.06 (0.06) | 0.00 | 0.20 | 0.06 (0.05) | 0.00 | 0.19 |
|  | *a* | 0.09 (0.08) | -0.08 | 0.25 | 0.09 (0.08) | -0.07 | 0.26 |
|  | $b_{2}$ | 0.06 (0.11) | -0.15 | 0.27 | 0.07 (0.10) | -0.13 | 0.27 |
| **POP3** | $\sigma_{proc}^{^{2}}$ | 0.23 (0.93) | 0.00 | 1.26 | 0.43 (1.58) | 0.00 | 2.41 |
|  | $\sigma_{obs}^{^{2}}$ | 0.11 (0.38) | 0.00 | 0.62 | 0.21 (0.94) | 0.00 | 1.13 |
|  | *a* | 0.36 (0.19) | 0.00 | 0.73 | 0.42 (0.27) | -0.09 | 0.93 |
|  | $b_{2}$ | -0.34 (0.25) | -0.81 | 0.13 | -0.24 (0.34) | -0.87 | 0.42 |
| **POP4** | $\sigma_{proc}^{^{2}}$ | 3.59 (3.37) | 0.04 | 9.89 | 2.53 (2.96) | 0.01 | 9.99 |
|  | $\sigma_{obs}^{^{2}}$ | 2.06 (1.99) | 0.01 | 7.07 | 2.40 (1.95) | 0.03 | 7.17 |
|  | *a* | 0.05 (0.50) | -1.00 | 1.09 | 0.03 (0.43) | -0.87 | 0.94 |
|  | $b_{2}$ | 0.06 (0.60) | -1.12 | 1.29 | -0.56 (0.69) | -1.91 | 0.82 |
| **POP5** | $\sigma_{proc}^{^{2}}$ | 0.23 (0.21) | 0.00 | 0.73 | 0.24 (0.21) | 0.00 | 0.74 |
|  | $\sigma_{obs}^{^{2}}$ | 0.15 (0.14) | 0.00 | 0.48 | 0.16 (0.14) | 0.00 | 0.49 |
|  | *a* | 0.09 (0.11) | -0.14 | 0.34 | 0.09 (0.11) | -0.14 | 0.34 |
|  | $b_{2}$ | 0.13 (0.16) | -0.18 | 0.45 | -0.09 (0.16) | -0.40 | 0.23 |

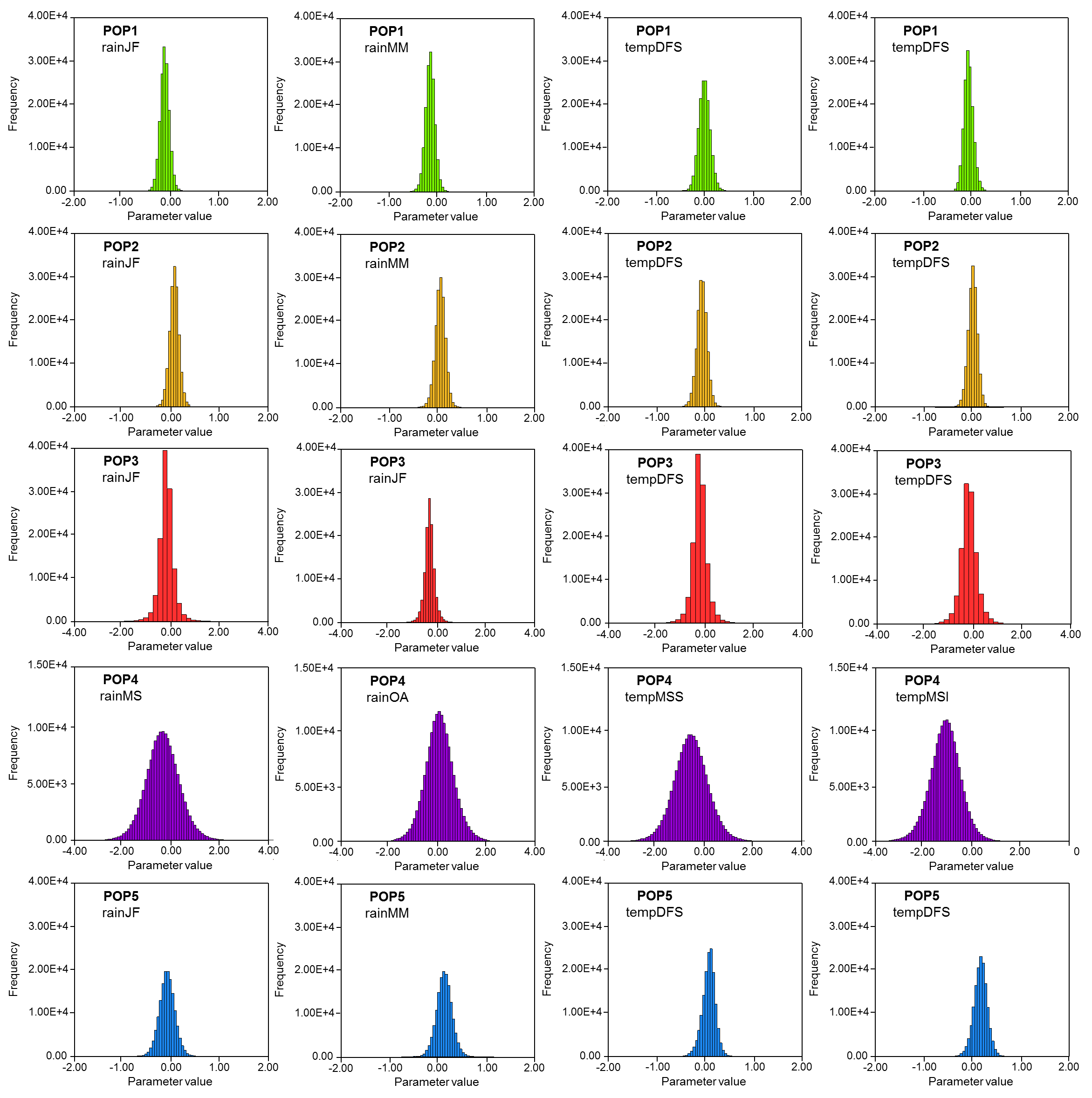

**Fig. S5.2.** Effect of local weather on the population growth rate in five populations of *Triturus cristatus* in Western Europe. The figure shows the distribution of posterior estimates for each of the four weather variables considered in each population.

As recapture probability may over time in several populations (Appendix 3), we examined if using population sizes corrected by time-dependent recapture probability (Horvitz-Thompson estimator with time-specific recapture) affected the detection of relationships between population growth and density and weather factors. We performed the same analyses as those presented above and reported the slope coefficient and the significance of the density and weather effects (**Fig.S5.2**). We showed that using Horvitz-Thompson estimator with time-specific or time-constant recapture probabilities weakly affects our results.

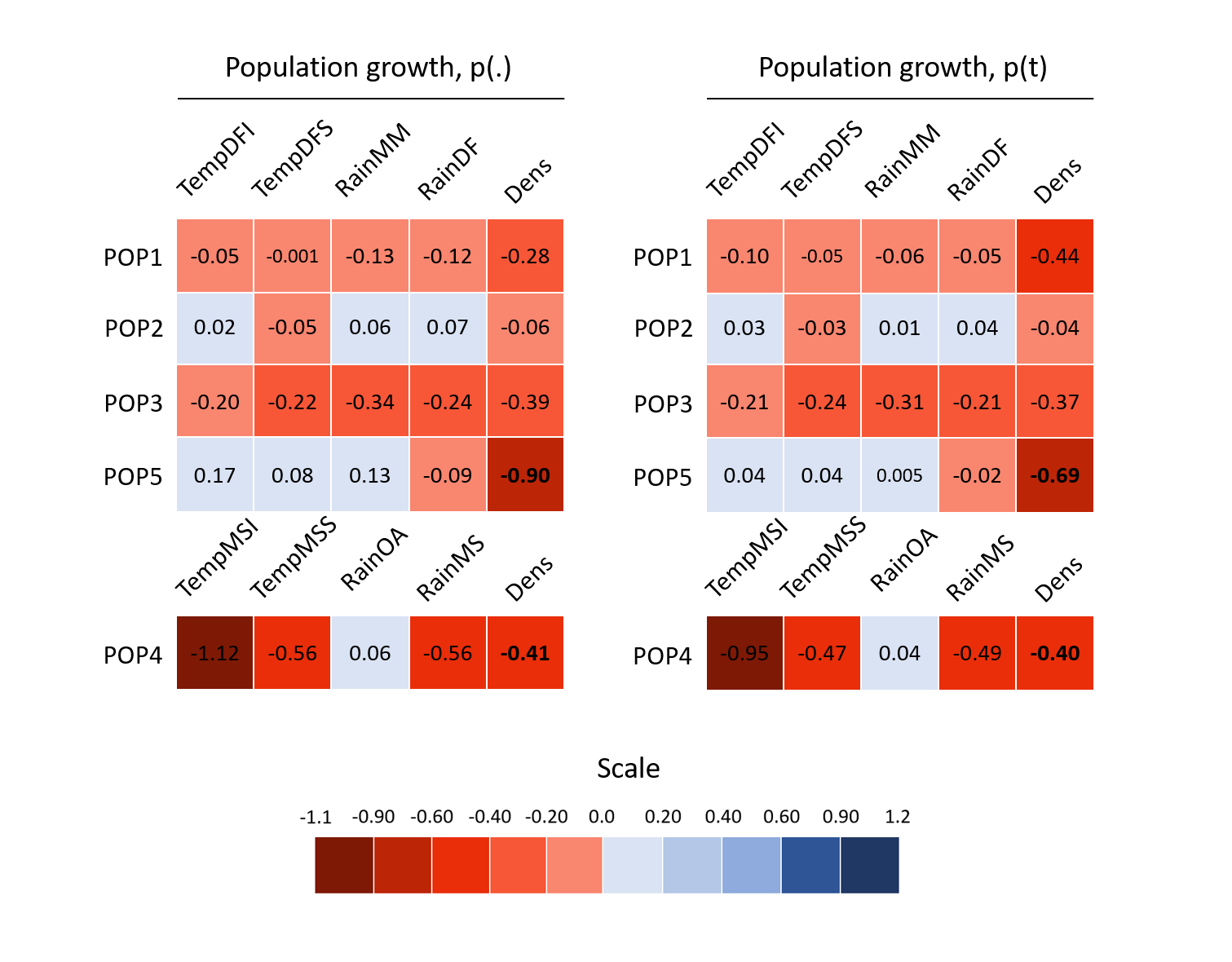

**Fig. S5.3.** Slope coefficient and significance of the effects of density and weather on population growth. The slope coefficient appears in bold when relationship is significant.

2. *Examining demographic growth rate at the subpopulation level*

**Table S5.4.** Effect of population density at *t* – 1 on the population growth rate in five populations of *Triturus cristatus* in Western Europe. The table shows the estimates (with their standard deviation in brackets) and 95% credible intervals of the parameters for the GSS model. $\sigma_{obs}^{^{2}}$ = the observation variance, $\sigma_{proc}^{^{2}}$ = the process variance, *a* = intrinsic rate of increase, $b_{1}$ = the coefficient slope for density at *t* – 1.

|  | Parameter | Estimate (sd) | 2.5% | 97.5% |
| --- | --- | --- | --- | --- |
| **POP1.1** | $\sigma_{proc}^{^{2}}$ | 0.21 (0.12) | 0.01 | 0.50 |
|  | $\sigma_{obs}^{^{2}}$ | 0.07 (0.11) | 0.00 | 0.38 |
|  | *a* | 1.80 (1.30) | -0.25 | 5.10 |
|  | $b_{1}$ | -0.41 (0.29) | -1.16 | 0.05 |
| **POP1.2** | $\sigma_{proc}^{^{2}}$ | 0.22 (0.23) | 0.00 | 0.80 |
|  | $\sigma_{obs}^{^{2}}$ | 0.31 (0.23) | 0.00 | 0.84 |
|  | *a* | 4.80 (1.96) | 1.12 | 8.89 |
|  | $b_{1}$ | -1.06 (0.43) | -1.92 | -0.25 |
| **POP1.3** | $\sigma_{proc}^{^{2}}$ | 0.47 (0.27) | 0.08 | 1.12 |
|  | $\sigma_{obs}^{^{2}}$ | 0.11 (0.15) | 0.00 | 0.51 |
|  | *a* | -0.06 (0.23) | -0.51 | 0.42 |
|  | $b_{1}$ | -0.12 (0.08) | -0.30 | 0.07 |
| **POP1.4** | $\sigma_{proc}^{^{2}}$ | 0.90 (0.51 | 0.12 | 2.12 |
|  | $\sigma_{obs}^{^{2}}$ | 0.30 (0.43) | 0.00 | 1.53 |
|  | *a* | 0.48 (0.42) | -0.25 | 1.42 |
|  | $b_{1}$ | -0.39 (0.28) | -1.06 | 0.07 |
| **POP5.1** | $\sigma_{proc}^{^{2}}$ | 1.53 (0.97) | 0.03 | 3.81 |
|  | $\sigma_{obs}^{^{2}}$ | 0.70 (0.92) | 0.00 | 3.22 |
|  | *a* | 1.00 (0.60) | 0.00 | 2.40 |
|  | $b_{1}$ | -0.49 (0.29) | -1.24 | -0.45 |
| **POP5.2** | $\sigma_{proc}^{^{2}}$ | 0.56 (0.45) | 0.02 | 1.61 |
|  | $\sigma_{obs}^{^{2}}$ | 0.38 (0.39) | 0.01 | 1.37 |
|  | *a* | 1.21 (0.78) | 0.03 | 3.05 |
|  | $b_{1}$ | -0.36 (0.18) | -0.98 | -0.02 |
| **POP5.3** | $\sigma_{proc}^{^{2}}$ | 0.19 (0.21) | 0.00 | 0.71 |
|  | $\sigma_{obs}^{^{2}}$ | 0.22 (0.19) | 0.00 | 0.69 |
|  | *a* | 2.34 (0.87) | 0.91 | 4.33 |
|  | $b_{1}$ | -0.56 (0.22) | -1.07 | -0.20 |
| **POP5.4** | $\sigma_{proc}^{^{2}}$ | 0.11 (0.12) | 0.00 | 0.40 |
|  | $\sigma_{obs}^{^{2}}$ | 0.16 (0.12) | 0.00 | 0.43 |
|  | *A* | 4.09 (1.65) | 1.15 | 7.57 |
|  | $b_{1}$ | -0.96 (0.39) | -1.78 | -0.27 |

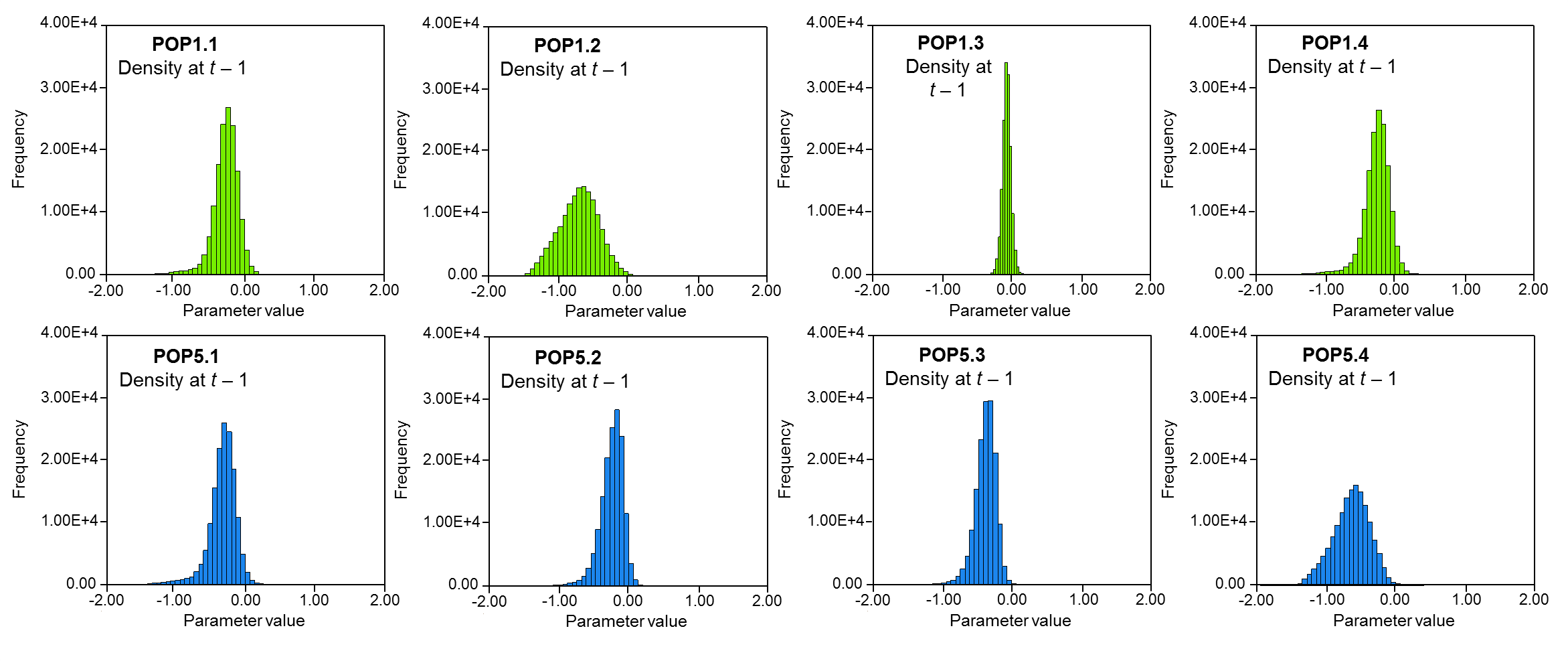

**Fig. S5.4.** Effect of population density at *t* – 1 on the population growth rate in the four subpopulations of POP1 and POP5. The figure shows the distribution of posterior estimates for density effects at *t* – 1.

**Table S5.5.** Effects of temperature on the population growth rate in five populations of *Triturus cristatus* in Western Europe. The table shows the estimates (with their standard deviation in brackets) and 95% credible intervals of the parameters for the GSS models including the local minimum and maximum temperature. $\sigma_{obs}^{^{2}}$ = the observation variance, $\sigma_{proc}^{^{2}}$ = the process variance, *a* = intrinsic rate of increase, $b_{2}$ = le slope for the temperature effect.

|  |  | Min. temperature  (tempDFI) | | | Max. temperature  (tempDFS) | | |
| --- | --- | --- | --- | --- | --- | --- | --- |
| Populations | Parameter | Estimate (sd) | 2.5% | 97.5% | Estimate (sd) | 2.5% | 97.5% |
| **POP1.1** | $\sigma_{proc}^{^{2}}$ | 0.23 (0.14) | 0.03 | 0.57 | 0.22 (0.14) | 0.01 | 0.55 |
|  | $\sigma_{obs}^{^{2}}$ | 0.06 (0.07) | 0.00 | 0.26 | 0.06 (0.08) | 0.00 | 0.29 |
|  | *a* | -0.01 (0.11) | -0.24 | 0.22 | -0.02 (0.11) | -0.24 | 0.21 |
|  | $b_{2}$ | -0.11 (0.13) | -0.36 | 0.14 | 0.13 (0.14) | -0.40 | 0.15 |
| **POP1.2** | $\sigma_{proc}^{^{2}}$ | 0.28 (0.34) | 0.00 | 1.20 | 0.23 (0.30) | 0.00 | 1.07 |
|  | $\sigma_{obs}^{^{2}}$ | 0.34 (0.23) | 0.01 | 0.87 | 0.34 (0.21) | 0.01 | 0.84 |
|  | *a* | -0.02 (0.13) | -0.29 | 0.26 | -0.02 (0.12) | -0.26 | 0.25 |
|  | $b_{2}$ | 0.01 (0.17) | -0.33 | 0.36 | 0.16 (0.18) | -0.20 | 0.53 |
| **POP1.3** | $\sigma_{proc}^{^{2}}$ | 0.51 (0.29) | 0.12 | 1.21 | 0.52 (0.29) | 0.11 | 1.21 |
|  | $\sigma_{obs}^{^{2}}$ | 0.11 (0.14) | 0.00 | 0.48 | 0.11 (0.15) | 0.00 | 0.52 |
|  | *a* | -0.25 (0.17) | -0.59 | 0.09 | -0.25 (0.17) | -0.59 | 0.10 |
|  | $b_{2}$ | -0.08 (0.19) | -0.45 | 0.30 | -0.05 (0.21) | -0.46 | 0.36 |
| **POP1.4** | $\sigma_{proc}^{^{2}}$ | 0.99 (0.61) | 0.17 | 2.48 | 1.01 (0.58) | 0.23 | 2.42 |
|  | $\sigma_{obs}^{^{2}}$ | 0.27 (0.32) | 0.00 | 1.13 | 0.23 (0.29) | 0.00 | 1.02 |
|  | *a* | 0.00 (0.24) | -0.48 | 0.48 | 0.01 (0.24) | -0.47 | 0.49 |
|  | $b_{2}$ | -0.05 (0.27) | -0.57 | 0.51 | 0.17 (0.29) | -0.40 | 0.75 |
| **POP5.1** | $\sigma_{proc}^{^{2}}$ | 1.72 (1.16) | 0.04 | 4.45 | 1.94 (1.21) | 0.07 | 4.78 |
|  | $\sigma_{obs}^{^{2}}$ | 0.63 (0.72) | 0.00 | 2.53 | 0.55 (0.70) | 0.00 | 2.45 |
|  | *a* | 0.16 (0.30) | -0.46 | 0.79 | 0.17 (0.32) | -0.49 | 0.84 |
|  | $b_{2}$ | 0.37 (0.37) | -0.38 | 1.10 | 0.19 (0.37) | -0.57 | 0.91 |
| **POP5.2** | $\sigma_{proc}^{^{2}}$ | 0.45 (0.50) | 0.00 | 1.74 | 0.65 (0.53) | 0.00 | 1.91 |
|  | $\sigma_{obs}^{^{2}}$ | 0.45 (0.33) | 0.00 | 1.23 | 0.34 (0.33) | 0.00 | 1.16 |
|  | *a* | 0.15 (0.16) | -0.19 | 0.49 | 0.14 (0.19) | -0.25 | 0.54 |
|  | $b_{2}$ | 0.14 (0.24) | -0.38 | 0.58 | -0.08 (0.24) | -0.60 | 0.36 |
| **POP5.3** | $\sigma_{proc}^{^{2}}$ | 0.52 (0.31) | 0.10 | 1.29 | 0.59 (0.33) | 0.12 | 1.39 |
|  | $\sigma_{obs}^{^{2}}$ | 0.15 (0.17) | 0.00 | 0.60 | 0.13 (0.17) | 0.00 | 0.57 |
|  | *a* | 0.24 (0.17) | -0.10 | 0.57 | 0.24 (0.18) | -0.13 | 0.59 |
|  | $b_{2}$ | 0.19 (0.21) | -0.22 | 0.59 | 0.03 (0.21) | -0.39 | 0.44 |
| **POP5.4** | $\sigma_{proc}^{^{2}}$ | 0.09 (0.14) | 0.00 | 0.49 | 0.12 (0.16) | 0.00 | 0.58 |
|  | $\sigma_{obs}^{^{2}}$ | 0.18 (0.10) | 0.01 | 0.43 | 0.18 (0.11) | 0.00 | 0.45 |
|  | *a* | 0.04 (0.07) | -0.11 | 0.20 | 0.05 (0.08) | -0.14 | 0.23 |
|  | $b_{2}$ | 0.16 (0.12) | -0.09 | 0.39 | 0.09 (0.11) | -0.15 | 0.30 |

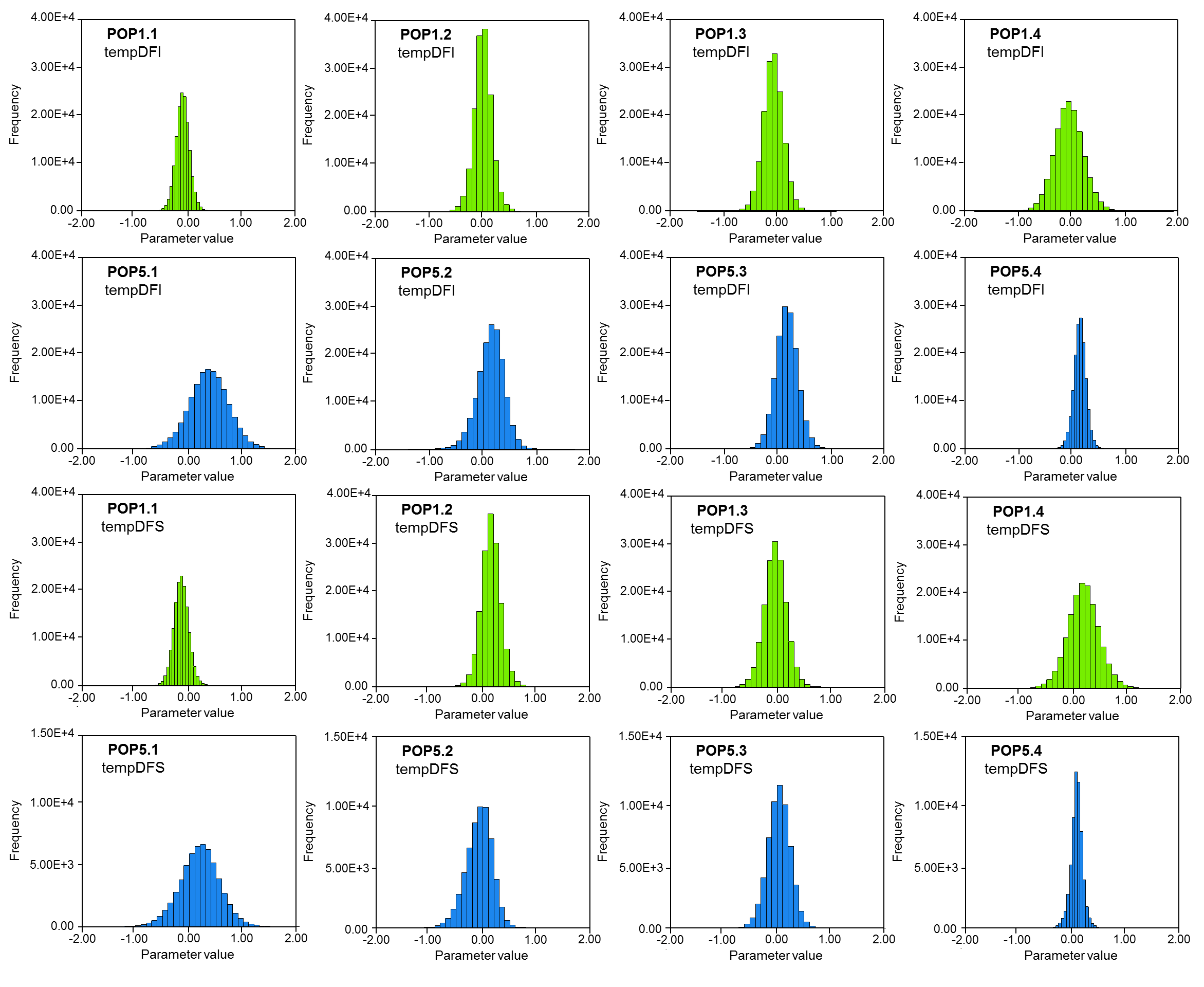

**Fig. S5.5.** Effect of local weather (tempDFI and tempDFS) on the population growth rate in the four subpopulations of POP1 and POP5. The figure shows the distribution of posterior estimates for the two weather variables

**Table S5.6.** Effect of rainfall on the population growth rate in five populations of *Triturus cristatus* in Western Europe. The table shows the estimates (with their standard deviation in brackets) and 95% credible intervals of the parameters for the GSS models including the local cumulative rainfall. $\sigma_{obs}^{^{2}}$ = the observation variance, $\sigma_{proc}^{^{2}}$ = the process variance, *a* = intrinsic rate of increase, $b_{2}$ = le slope for the rainfall effect.

|  |  | Rainfall (rainMM) | | | Rainfall (rainJF) | | |
| --- | --- | --- | --- | --- | --- | --- | --- |
| Populations | Parameter | Estimate (sd) | 2.5% | 97.5% | Estimate (sd) | 2.5% | 97.5% |
| **POP1.1** | $\sigma_{proc}^{^{2}}$ | 0.21 (0.13) | 0.03 | 0.53 | 0.24 (0.14) | 0.03 | 0.58 |
|  | $\sigma_{obs}^{^{2}}$ | 0.06 (0.07) | 0.00 | 0.26 | 0.06 (0.08) | 0.00 | 0.27 |
|  | *a* | -0.01 (0.11) | -0.23 | 0.21 | -0.01 (0.12) | -0.24 | 0.23 |
|  | $b_{2}$ | -0.14 (0.13) | -0.40 | 0.12 | -0.05 (0.14) | -0.32 | 0.22 |
| **POP1.2** | $\sigma_{proc}^{^{2}}$ | 0.25 (0.32) | 0.00 | 1.13 | 0.25 (0.30) | 0.00 | 1.06 |
|  | $\sigma_{obs}^{^{2}}$ | 0.35 (0.23) | 0.01 | 0.89 | 0.30 (0.20) | 0.00 | 0.79 |
|  | *a* | -0.01 (0.12) | -0.27 | 0.25 | -0.02 (0.12) | -0.09 | 0.25 |
|  | $b_{2}$ | -0.10 (0.19) | -0.48 | 0.27 | -0.21 (0.18) | -0.27 | 0.13 |
| **POP1.3** | $\sigma_{proc}^{^{2}}$ | 0.50 (0.27) | 0.13 | 1.17 | 0.42 (0.22) | 0.12 | 0.98 |
|  | $\sigma_{obs}^{^{2}}$ | 0.10 (0.13) | 0.00 | 0.44 | 0.08 (0.10) | 0.00 | 0.35 |
|  | *a* | -0.25 (0.17) | -0.59 | 0.08 | -0.25 (0.15) | -0.55 | 0.06 |
|  | $b_{2}$ | -0.15 (0.19) | -0.52 | 0.23 | -0.30 (0.17) | -0.64 | 0.04 |
| **POP1.4** | $\sigma_{proc}^{^{2}}$ | 0.65 (0.42) | 0.07 | 1.68 | 0.91 (0.54) | 0.18 | 2.42 |
|  | $\sigma_{obs}^{^{2}}$ | 0.20 (0.24) | 0.00 | 0.85 | 0.23 (0.28) | 0.00 | 1.02 |
|  | *a* | 0.00 (0.19) | -0.40 | 0.39 | 0.00 (0.23) | -0.46 | 0.49 |
|  | $b_{2}$ | -0.55 (0.23) | -0.99 | -0.09 | -0.30 (0.26) | -0.81 | 0.21 |
| **POP5.1** | $\sigma_{proc}^{^{2}}$ | 2.03 (1.18) | 0.17 | 4.81 | 2.00 (1.15) | 0.27 | 4.74 |
|  | $\sigma_{obs}^{^{2}}$ | 0.51 (0.69) | 0.00 | 2.42 | 0.50 (0.64) | 0.00 | 2.28 |
|  | *a* | 0.16 (0.33) | -0.50 | 0.84 | 0.14 (0.33) | -0.51 | 0.80 |
|  | $b_{2}$ | 0.17 (0.41) | -0.64 | 0.98 | 0.23 (0.40) | -0.56 | 1.03 |
| **POP5.2** | $\sigma_{proc}^{^{2}}$ | 0.61 (0.51) | 0.00 | 1.83 | 0.57 (0.52) | 0.00 | 1.85 |
|  | $\sigma_{obs}^{^{2}}$ | 0.37 (0.34) | 0.00 | 1.20 | 0.41 (0.35) | 0.00 | 1.24 |
|  | *a* | 0.13 (0.18) | -0.25 | 0.52 | -0.14 (0.18) | -0.23 | 0.52 |
|  | $b_{2}$ | -0.12 (0.28) | -0.67 | 0.42 | 0.00 (0.26) | -0.50 | 0.52 |
| **POP5.3** | $\sigma_{proc}^{^{2}}$ | 0.56 (0.32) | 0.11 | 1.34 | 0.58 (0.32) | 0.14 | 1.36 |
|  | $\sigma_{obs}^{^{2}}$ | 0.14 (0.17) | 0.00 | 0.60 | 0.14 (0.17) | 0.00 | 0.57 |
|  | *a* | 0.24 (0.17) | -0.11 | 0.59 | 0.23 (0.18) | -0.12 | 0.59 |
|  | $b_{2}$ | 0.15 (0.22) | -0.28 | 0.59 | -0.09 (0.22) | -0.53 | 0.34 |
| **POP5.4** | $\sigma_{proc}^{^{2}}$ | 0.14 (0.17) | 0.00 | 0.58 | 0.15 (0.17) | 0.00 | 0.60 |
|  | $\sigma_{obs}^{^{2}}$ | 0.16 (0.11) | 0.00 | 0.43 | 0.17 (0.12) | 0.00 | 0.45 |
|  | *a* | 0.04 (0.09) | -0.14 | 0.25 | 0.03 (0.09) | -0.15 | 0.24 |
|  | $b_{2}$ | 0.17 (0.14) | -0.11 | 0.47 | -0.11 (0.14) | -0.41 | 0.17 |

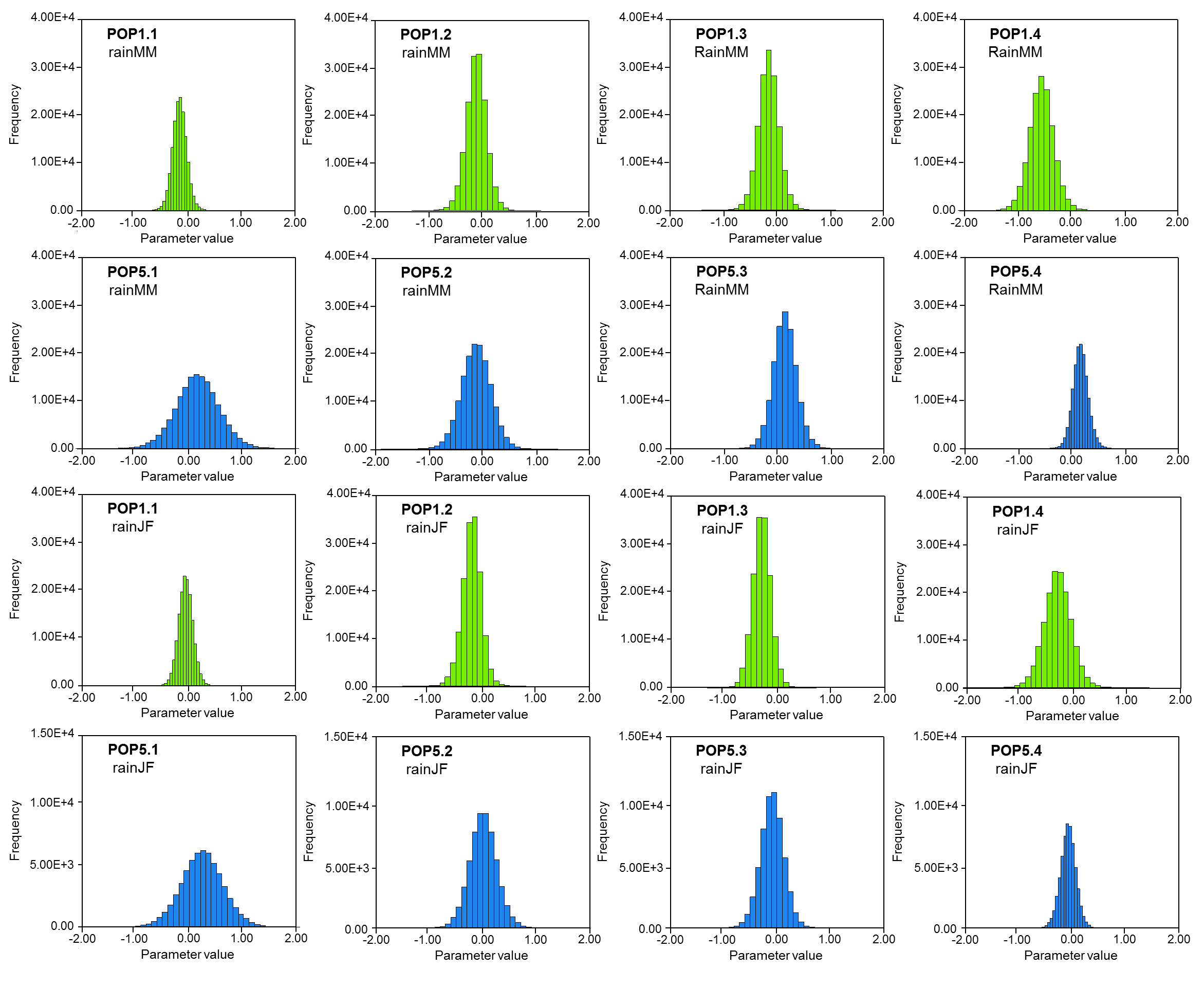

**Fig. S5.6.** Effect of local weather (rainMM and rainJF) on the population growth rate in the four subpopulations of POP1 and POP5. The figure shows the distribution of posterior estimates for the two weather variables.
